# Mechanistic Explanation of Neuroplasticity Using Equivalent Circuits

**DOI:** 10.1101/2023.05.21.541639

**Authors:** Martin N. P. Nilsson

**Author notes:** The author declares no competing financial interests.

## Abstract

This paper presents a comprehensive mechanistic model of a neuron with plasticity that explains how information input as time-varying signals is processed and stored. Additionally, the model addresses two long-standing, specific biological challenges: Integrating Hebbian and homeostatic plasticity, and identifying a concise synaptic learning rule.

A biologically accurate electric-circuit equivalent is derived through a one-to-one mapping from the known properties of ion channels. The often-overlooked dynamics of the synaptic cleft is essential in this process. Analysis of the model reveals a simple and succinct learning rule, indicating that the neuron functions as an internal-feedback adaptive filter, a common concept in signal processing. Simulations confirm the model’s functionality, stability, and convergence, demonstrating that even a single neuron without external feedback can act as a potent signal processor.

The model replicates several key characteristics typical of biological neurons, which are seldom captured in other neuron models. It can encode time-varying functions, learn without risking instability, and bootstrap from a state where all synaptic weights are zero.

This paper explores the function of neurons with a focus on biological accuracy, not computational efficiency. Unlike neuromorphic models, it does not aim to design devices. The electronic circuit analogy aids understanding by leveraging decades of electronics expertise but is not intended for physical implementation.

This interdisciplinary work spans a broad range of subjects within the realm of neurobiophysics, including neurobiology, electronics, and signal processing.

**Significance statement:** Mechanistic neuron models with plasticity are crucial for understanding the complexities of the brain and the processes behind learning and memory. These models provide a way to study how individual neurons and synapses in the brain change over time in response to stimuli, allowing for a more nuanced understanding of neuronal circuits and assemblies. Plasticity is a key aspect of these models, as it represents the ability of the brain to modify its connections and functions in response to experiences. By incorporating plasticity into these models, researchers can explore how changes at the synaptic level contribute to higher-level changes in behavior and cognition. Thus, these models are essential for advancing our understanding of the brain and its functions.

**PhySH terms:** Neuroplasticity, Neural basis of learning and memory, Synapses.

**MeSH 2023 terms:** Neuronal Plasticity [G11.561.638], Association learning [F02.463.425.069.296], Memory [F02.463.425.540], Synaptic transmission [G02.111.820.850]

## Introduction

How does the brain remember? This classical question has recently received considerable attention focusing on the central nervous system’s handling of co-ordinated synaptic changes, mandating a cell-wide coherent explanation of multisynaptic plasticity. However, a mechanistic explanation remains to be found, despite massive research efforts by experimental and theoretical methods. This paper suggests that the question can be answered by mapping the neuron to an equivalent circuit and then showing that this circuit implements an adaptive filter. While many researchers have proposed that neurons implement such filters (*e*.*g*., [76, 18, 78, 57, 42]), to the best of the author’s knowledge, no one has yet provided an explanation of how the filter function is achieved on the ion channel level.

Experimental methods typically investigate neurons’ responses to stimuli and various biological manipulations such as ion channel blocking and genetic modifications. Since the first discovery of synaptic plasticity [5], experiments have revealed a diversity of overlapping and interacting plasticity mechanisms [40, 68]. A limitation of experiments *in vitro* is that crucial parameters such as temperature, membrane potential, and calcium concentration often transcend their physiological ranges. On the other hand, experiments *in vivo* are degraded by external disturbances, such as irrelevant signals from connected neurons.

The theoretical approach is to study how plasticity *should* work. In this case, the principal obstacle is to find biologically plausible mechanisms that match the theory. A significant theoretical contribution showed that classical Hebbian plasticity alone leads to the saturation of synaptic weights and the ensuing loss of information [53, 3]. Subsequent experiments demonstrated the existence of additional, homeostatic mechanisms that prevent distortion and stabilize synaptic plasticity [72, 71]. Here, “homeostasis” refers to the neuron’s retaining a stable internal environment despite changes in external conditions.

Because of the challenges posed by the diversity of plasticity mechanisms and the scarcity of biologically plausible models, the timing and integration of homeostatic and Hebbian plasticity is an open issue [34]. Therefore, a novel approach is chosen here, modeling a neuron as an electric-circuit equivalent in the spirit of Hodgkin and Huxley’s seminal model [29] while strictly adhering to known properties of neuronal ion channels to ensure biological veracity. It should be noted that the Hodgkin-Huxley model is a model of a neuron’s *axon*—specifically, the giant axon of *Loligo*. As such, it describes the *output* section of the neuron, *i*.*e*., how the neuron converts membrane potential to spike trains. In contrast, the current paper describes how the *input* section converts spike trains back to membrane potential, including an explanation of plasticity.

The original Hodgkin-Huxley model is a deterministic, single-compartment model that accurately predicts the typical shape of neuronal spikes but falls short in explaining the stochastic variability of interspike intervals (ISI). However, the ISI probability distribution can be accurately modeled by dividing the single compartment into three distinct compartments, consisting of the distal compartment, which includes the distal dendrites; the proximal compartment, housing the proximal dendrites and the soma; and the axon initial segment (AIS) [49]. Notably, for the analysis of the output section it is unnecessary to incorporate any considerations of synapses or plasticity. The principal mathematical difficulty of the *output* section—the proximal compartment and the AIS—lies in solving a small number of stochastic differential equations. This is undertaken in [49], which employs the Cramér-Rao lower bound to show that the model cannot be significantly improved unless experimental data are vastly improved.

The challenges involved in analyzing the *input* section are distinctly different. A single excitatory synapse in the model is represented by 32 coupled equations, including four non-linear and five differential equations (cf. appendix). As a result, classical analysis methods become inadequate due to the excessive complexity when modeling a complete neuron. To address this, the paper leverages insights from electronics. The resulting circuit can be interpreted mechanistically as a modified *Least Mean Square* (LMS) *adaptive filter*, a versatile device well-known in the field of signal processing [27, 26]. This interpretation takes advantage of the rich theory developed for adaptive filters. It explains precisely and quantitatively how the neuron modifies its synapses in orchestration to store time-variable functions or *signals* as required by procedural memory.

Many previous attempts to explain plasticity focus on AMPA (*α*-amino-3-hydroxy-5-methyl-4-isoxazolepropionic acid) receptors, assuming they not only carry the primary feed-forward signal but also control plasticity. However, this assumption leads to complications. Silent synapses, where the synaptic weight is zero, cannot convey any signal, which would cause the synapse to remain stuck at zero. Even if we assume that the synaptic weight is slightly positive, problems arise because the charge entering an AMPA receptor is directly summed into the membrane potential, making the local synapse’s contribution indistinguishable from other inputs.

The above problems indicate that an additional channel appropriately reflects pre-synaptic activity, with calcium entry via NMDA (N-methyl-D-aspartate) receptors appearing to be the most viable candidate [31, 70, 51, 44]. Whereas it is well known that variations in external calcium concentration are minuscule, the calcium flow variations being small does not pose issues in the proposed model other than slowing down adaptation because the model behaves linearly and remains stable for small excursions from equilibrium. Biological experiments often require exceptionally strong stimuli, such as tetanic stimulation, to detect short-term plasticity effects. Accordingly, the experiments described here intentionally use exaggerated calcium variations to make the adaptation process more conspicuous. A more profound reason for using large variations is that significant deviations in a feedback system carry the risk of instability and functional collapse. The presented experiments demonstrate that the circuit remains stable despite substantial deflections from equilibrium.

The importance of mechanistic models cannot be overstated, as these models offer significant benefits over empirical or phenomenological models. Their superiority stems from their ability to deliver more comprehensive and detailed insight into biological and physiological processes. Mechanistic models excel, particularly regarding

- **Explanation and causation:** Mechanistic models are designed to explain phenomena not merely through observation but by delineating the underlying mechanisms that cause these observations. While empirical models correlate inputs and outputs and phenomenological models describe relationships, mechanistic models explain how one state leads to another by interacting components within the system.
- **Predictive power:** By detailing the specific interactions and processes, mechanistic models often possess superior predictive capabilities, especially in conditions that have not been directly observed.
- **Innovation and hypothesis testing:** Mechanistic models allow scientists to test hypotheses about the effects of modifying system components. This can be invaluable for experimental biology, where understanding the role of a specific gene or protein in a pathway can lead to breakthroughs in treatment methods. Mechanistic models provide a framework within which new ideas can be tested theoretically before moving to costly and time-consuming experiments.
- **Integration of information:** These models integrate information across different levels of biological organization—from molecular to cellular to organismal levels—thereby allowing a holistic understanding of complex biological systems. For instance, a mechanistic model of neuron function can incorporate both molecular interactions at the synapse and the effect of these interactions on overall neuronal activity and network dynamics.
- **Education and communication:** Mechanistic models serve as practical educational tools that help new researchers and practitioners understand the complex interactions underpinning the systems they study. They also aid in communicating scientific concepts to non-experts or interdisciplinary teams, fostering broader collaboration.
- **Refinement of theory:** As new data become available, mechanistic models can be refined and expanded. This iterative process helps continuously improve the accuracy and relevance of scientific theories, ensuring that they remain robust in the face of new evidence.

Given these points, the reliance on a mechanistic approach in the proposed model—as investigated in [10] and [11]—is essential not just for accuracy and depth of understanding, but also for the practical implications it has in research and application. Mechanistic models bridge the gap between theory and practice, making them indispensable tools in the advancement of science.

It is tempting to include numerous features of ion channels in the model with hopes of the plasticity function emerging from the model. However, such an approach is unlikely to succeed, as it would result in excessive complexity. Additionally, many ion channels serve disparate functions such as homeostasis, metabolism, or potentially, immune system regulation. This would make it challenging to discern which ion channel features are relevant to the function being studied and when the model is sufficiently comprehensive.

Instead, the approach advocated here aligns with Occam’s razor principle: the model should be as simple as possible while including established features that adequately explain the phenomenon under consideration. Newton’s observation of an apple falling from a tree serves as a famous example. He didn’t deem it necessary to incorporate air resistance or buoyancy into his model, opting for highly simplified representations of the earth and the apple. Similarly, we adhere to this principle by limiting the model to well-established features of ion channels, incorporating them only when they contribute significantly to explaining neuroplasticity.

The purpose of this paper is to explain how biological neurons function, with a focus on biological accuracy. Here, computational efficiency is irrelevant. This distinction is important, as it differs from the goals of neuromorphic models, which use biological inspiration to design efficient computational devices without requiring biological accuracy. The use of an electronic circuit analogy for the neuron should not be construed as an attempt to construct a physical device. Instead, it is a method that leverages decades of experience in electronics to facilitate our understanding of the neuron’s complex biophysical processes.

This paper is highly interdisciplinary, which presents challenges for readers from different academic backgrounds. Biologists are typically well-acquainted with the importance of mechanistic models but may be less familiar with signal processing principles, feedback systems, and related concepts rooted in classical engineering. In contrast, computational neuroscientists often rely on established neuron simulators and may lack exposure to foundational biological concepts such as Gray’s rules or the rationale behind mechanistic modeling.

Additionally, modeling in terms of active electronic components is uncommon in both fields, yet this perspective is essential for the present work. The biological, signal processing, and electronic aspects are all critical and cannot be excluded without compromising the integrity of the overall approach.

To support a broad readership, care has been taken to include sufficient detail across these domains. Naturally, some readers may find certain sections overly detailed, while others may consider the same material insufficient. Given space constraints, a balance has been sought. For those requiring additional background, the following baseline resources are recommended: [58, Ch. 1-8] for neurobiology, [30, Ch. 1-7] for electronics, and [77, Ch. 1-12] for adaptive signal processing.

### Organization of this paper

The paper’s main topic is a derivation of the equivalent circuit and adaptive filter model from established knowledge about neuronal ion channels. For a mechanistic model, it is imperative to select an appropriate level of description that is adequately detailed yet not overly complex to provide a functional explanation and address the three specific problems under consideration. To achieve this, the paper first reviews the established function of inhibitory and excitatory synaptic ion channels to a level that allows for a direct translation into an electric network. By this conversion, insights from a century of experience with electronic circuits can be leveraged, along with the ability to identify circuit patterns or “motifs.” The approach is conservative in not assuming the existence of as-yet-undiscovered biological mechanisms.

The paper’s main conclusion is that a single neuron can be abstractly and mechanistically characterized as an adaptive filter, a powerful and fundamental component in signal processing. The basic principles of adaptive filters are, therefore, briefly reviewed. An adaptive filter’s function in its fullest generality is to determine how a reference input is expressible in terms of a given set of input components.

Two experiments are performed to support further the claim that the neuron operates as an adaptive filter. The first experiment demonstrates the circuit’s ability to approximate an inhibitory signal *y*(*t*) by appropriately weighting excitatory inputs *x*_*k*_(*t*). The second experiment confirms that action potentials function as clock pulses (“strobes”), triggering synaptic weight changes.

The results section presents the convergence and stability outcomes of the experiments diagrammatically, followed by an explanation of how the model, in its adaptive filter capacity, addresses the three specific issues concerning memory, Hebbian-homeostatic plasticity, and the synaptic learning rule.

Subsequently, the discussion section introduces related work and explores some implications of viewing the neuron as an adaptive filter.

The investigation spans a time frame from milliseconds to minutes, encompassing short-term plasticity (STP) and early long-term potentiation (LTP) while excluding late LTP due to its reliance on nuclear processes and its consolidating function.

The neuron model introduced here lays the groundwork for a more complex mechanistic model that examines neuron populations and their coding mechanisms. However, creating a model encompassing large networks of neurons requires sophisticated signal processing techniques, such as wavelet decomposition and the concept of sparsity. These aspects are beyond the scope of the current paper but are detailed in a separate study [48]. To summarize that study briefly, it demonstrates how populations of neurons conforming to the adaptive-filter model discussed here can effectively transmit, process, and store information. This is achieved through an invariance property, which can be geometrically characterized as a convex cone. The adaptive filtering characteristics of these neurons enable them to perform signal processing tasks compactly and efficiently. An algebra of convex cones can abstractly describe these operations. This provides the populations with a robust computational framework akin to a “programming language” for neurons.

In summary, this article models a neuron’s primary biochemical information processing pathways as equivalent electric circuits, reviews the adaptive filter concept, and employs it to describe the neuron’s overall function. The model’s adequacy is demonstrated through two simulation experiments, substantiating the neuron’s capacity to operate as an adaptive filter. These results support the proposed model’s validity and potential for advancing research in this field, demonstrated by its application as a foundation for a mechanistic model of neuron populations [48].

## Modeling the neuron

### Overall structure of a neuron

This subsection provides a detailed description of the structure and function of a neuron, highlighting its key components, synaptic types, and their roles in signal transmission and plasticity.

The target neuron is a generic, glutamatergic neuron equipped with AMPA receptors (AMPARs) and NMDA receptors (NMDARs). This kind of neuron has been extensively studied and is representative of a substantial fraction of neurons in the central nervous system (CNS) [70], typical examples of which are the hippocampal neurons where plasticity was first demonstrated [5].

The primary components of a neuron include the dendrites, which receive inputs from presynaptic neurons; the soma, which aggregates the contributions from dendrites; and the axon, which transmits the result to other neurons (fig. 1). Axons can branch into axon collaterals carrying identical signals. Synapses, the contact points between axons and dendrites, are of two types: inhibitory and excitatory. They convert incoming stochastically rate-coded sequences of action potentials (APs), or more tersely, PFM (pulse-frequency modulated) spiketrains [49], into postsynaptic currents that alter the membrane potential, the voltage difference between the neuron’s interior and exterior. At the axon initial segment (AIS), this potential is converted back into a spiketrain for output via the axon.

**Figure 1:**
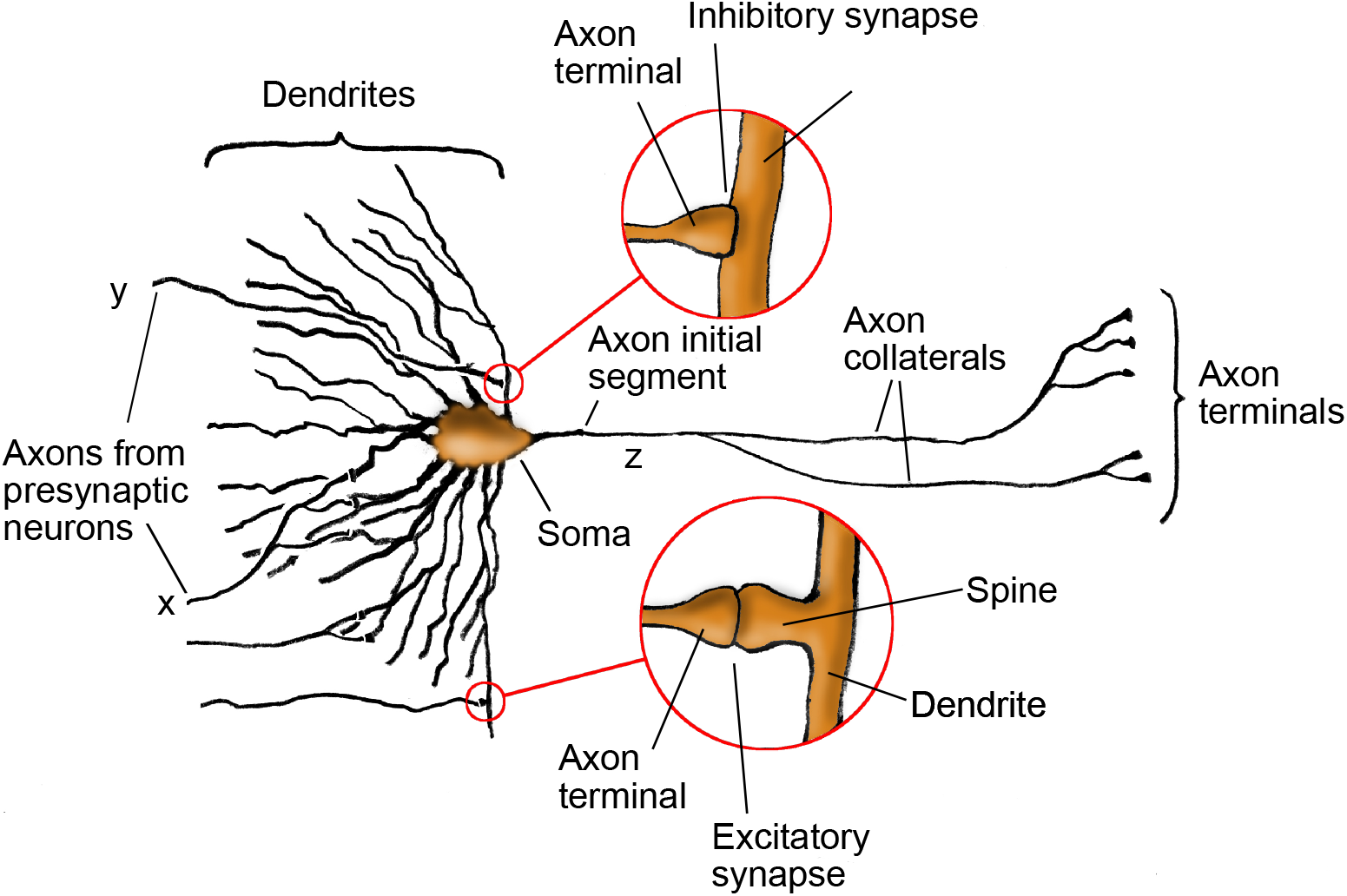
Structure of a generic neuron. This image overviews the essential parts of a neuron, with details of inhibitory and excitatory synapses explored in fig. 2 and fig. 5, respectively.

Ultrastructural studies reveal that inhibitory synapses are typically situated proximally to the soma, directly on a dendrite or the soma itself. In contrast, excitatory synapses tend to be positioned more distally, connecting with dendrites via spines—small protrusions on the dendrites. These spines are associated with plasticity, indicating that excitatory synapses are generally plastic, whereas inhibitory synapses are non-plastic. These principles are occasionally referred to as Gray’s rules [23, 52, 70, 25].

### Inhibitory synapse mapping

This subsection focuses on the functioning of an inhibitory synapse in response to an action potential, including the involvement of neurotransmitters and receptors. The subsection presents the corresponding equivalent electric-circuit model that reflects this process, inspired by Hodgkin and Huxley’s axon model [29].

The flow of neurotransmitters or charges, such as ions and electrons, are modeled as currents. In addition to representing themselves, voltages also depict accumulations and concentrations of ions. We define ground as the local resting potential *E*_rest_ and focus on slight deviations from this baseline by the membrane potential. Because the analysis focuses on these small-signal (AC) variations, the exact values of the voltage sources and reference voltages in the circuits are less crucial.

The model adopts Hodgkin and Huxley’s view of gated ion channels as voltage-controlled conductances (fig. 3A). Because of their similarity to ideal field-effect transistors (FETs), the schematic uses modern transistor symbols (fig. 3B, C). A formal rationale that a local population of gated ion channels can be identified as a transistor is that they are governed by the same constitutive equation,

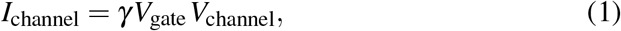

saying that the channel current is proportional to the product of the gate and channel voltages, where the factor *γV*_gate_ denotes the channel conductance.

When action potentials reach the axon terminal (fig. 2), the membrane potential depolarizes (increases), causing voltage-gated calcium channels (Ca_V_) to open (1). The calcium ion influx triggers the release of the neurotransmitter *γ*-aminobutyric acid (GABA) from nearby vesicles (2) into the synaptic cleft. GABA binds to GABA type A receptors (GABAAR) on the postsynaptic neuron, opening the receptor channel to chloride ions (3) [62]. These ions are negatively charged and hyperpolarize (reduce) the membrane potential. A direct translation of these biological processes into a circuit equivalent for the inhibitory synapse is shown in fig. 4.

**Figure 2:**
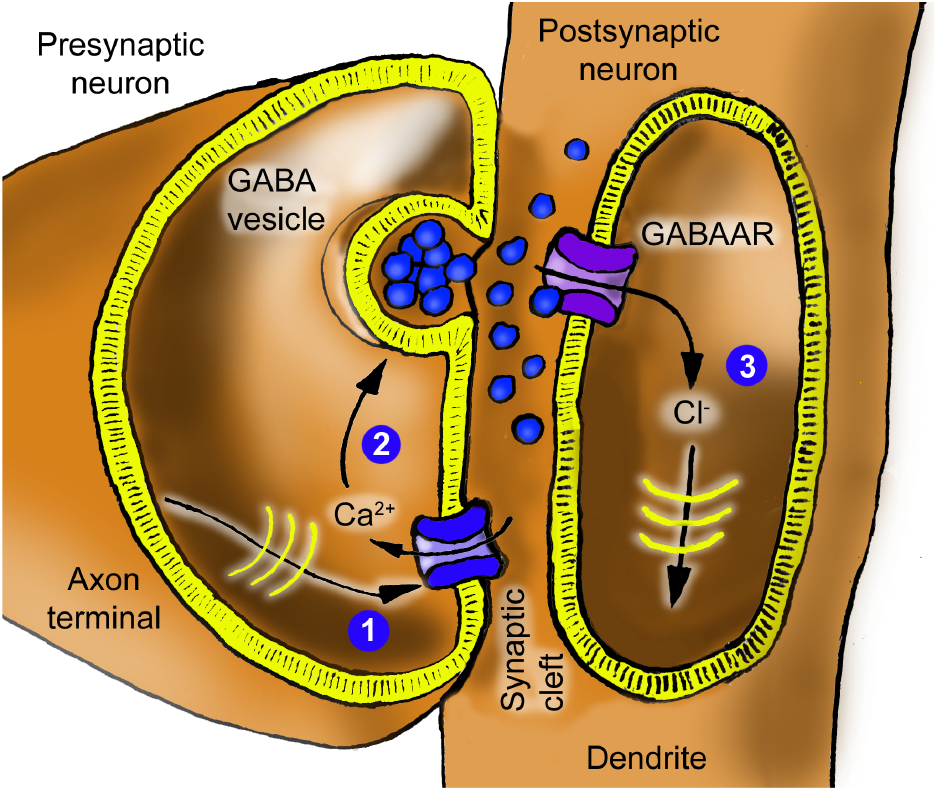
Inhibitory synapse schematic. An action potential arriving at (1) causes the release of GABA at (2), which then activates GABAAR, allowing a chloride current at (3), traveling as an IPSC to the soma.

**Figure 3:**
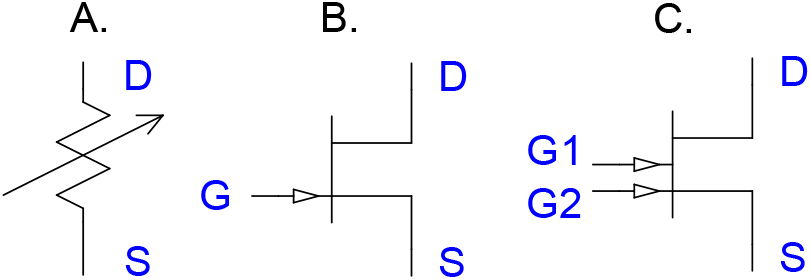
Gated ion channels are transistors. Gated ion channels are FET transistors. **A**. Hodgkin and Huxley used a variable resistance to describe gated ion channels. In their model, the gating connection was not drawn explicitly but appeared in the constitutive equation for the channel. **B**. Today, the transistor concept is well-established and, in particular, an ideal FET (field effect transistor) is an excellent model for a gated ion channel. A voltage-gated calcium channel Ca_V_ is accurately described by a transistor where the gate voltage *V*_*G*_ is provided by the membrane potential, controlling the drain-source (DS) current *I*_*DS*_ by the constitutive equation *I*_*DS*_ = *γV*_*G*_*V*_*DS*_. Similarly, a transistor can be used for the GABAAR, where the gate voltage models the GABA concentration. **C**. Double-gated transistors appropriately model NMDAR and AMPAR defined by *I*_*DS*_ = *γV*_*G*1_*V*_*G*2_*V*_*DS*_, involving the two gating inputs *G*1 and *G*2. For the NMDAR, these are the glutamate concentration and the postsynaptic membrane potential, respectively, whereas for the AMPAR, they are the glutamate concentration and the number of AMPAR receptors, respectively.

**Figure 4:**
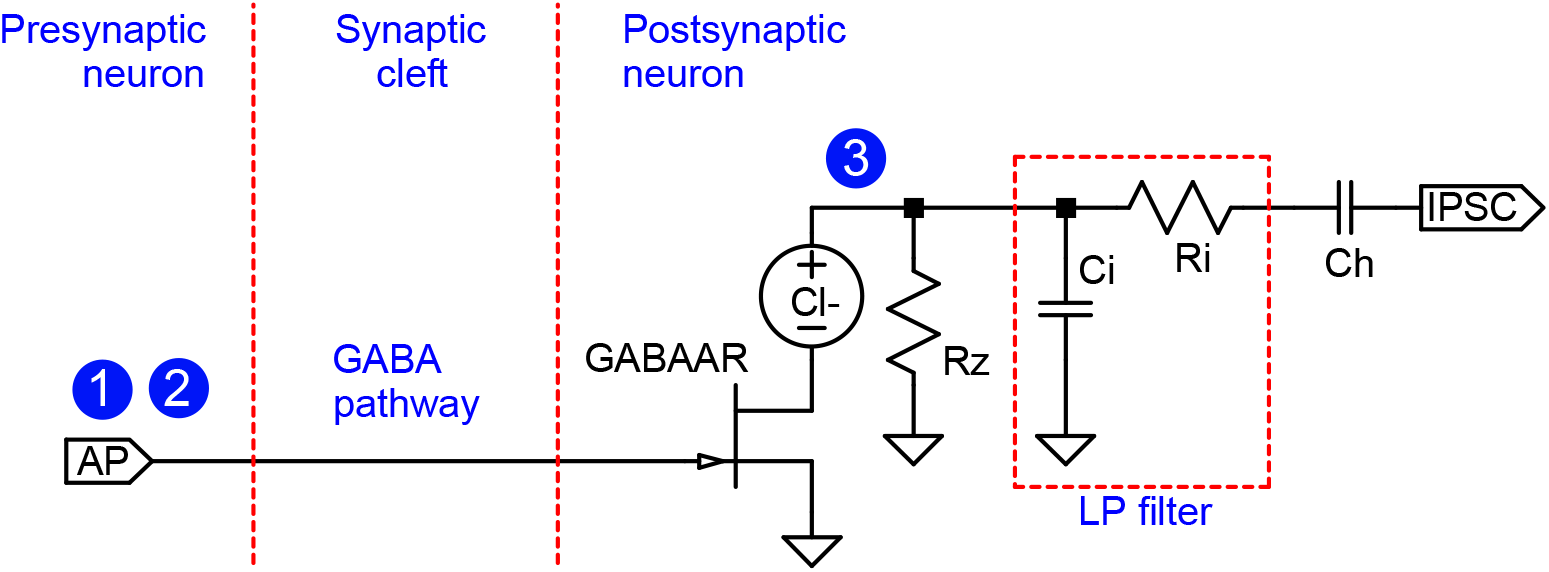
Inhibitory synapse circuit equivalent. This circuit implements the inhibitory synapse described in fig. 2 by modeling neurotransmitters as electrical currents. The transistor represents GABA-gated ion channels. Locations indicated by circled numbers 1-3 correspond to identically marked locations in fig. 2. As long as the membrane potential is higher than the reversal potential of chloride, which is normally the case, a positive pulse input at AP will lead to a negative current pulse at IPSC. The capacitor *C*_*h*_ models the cellular machinery retaining homeostatic equilibrium.

The GABAAR is defined by the equation *I*_*DS*_ = *γV*_*G*_*V*_*DS*_, where the constant *γ* = *γ*_GABAAR_ is the transistor’s gain, and *I*_*DS*_ and *V*_*DS*_ = *V*_*D*_ − *V*_*S*_ are the channel current and voltage, respectively, between the transistor’s *D* (“drain”) and *S* (“source”) terminals or “pins.” *V*_*G*_ is the gate voltage representing the GABA concentration. The voltage source “Cl-” represents the offset of the equilibrium potential *E*_rest_ from the reversal potential *E*_Cl_ for chloride ions, so

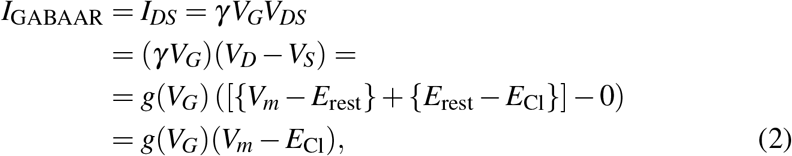

recognizable as the traditional equation for the ion channel current where *g*(*V*_*G*_) is the conductance, *V*_*m*_ is the local membrane potential at (3) in fig. 4, and *V*_*m*_ − E_Cl_ is the electrochemical driving force.

A single transistor is chosen to represent the entire population of GABAARs at one synapse. Overall, the circuit inverts an incoming train of positive voltage pulses to negative current pulses and filters them through a lowpass filter before integrating them into the membrane potential.

The resistor *R*_*z*_ represents the transport processes that circulate the chloride back out of the cell. The signal is filtered on its way to the soma by a lowpass filter *R*_*i*_*C*_*i*_ composed of the spino-dendritic axial resistance *R*_*i*_ and capacitance *C*_*i*_. The filter properties of synapses can vary depending on their proximity to the soma. In the current inhibitory synapse model, this variability can be represented by adjusting the *R*_*i*_ and *C*_*i*_ components. Nonetheless, to maintain simplicity in the explanation, this feature is not used in the example scenarios below.

The series capacitance *C*_*h*_ effectively encapsulates the homeostatic mechanisms that stabilize the neuron’s internal potential at a biologically optimal level. Alternatively, it can be viewed as a means of compartmentalizing the neuron’s functional regions. In a single-compartment neuron model, the membrane resistance *R*_*m*_ and membrane capacitance *C*_*m*_ largely determine the neuron’s frequency characteristics. However, as demonstrated in [49], a mechanistic explanation of spike generation requires at least three distinct compartments. In this framework, the resistor *R*_*z*_ serves to establish a local reference potential for the distal compartment and the *R*_*i*_ and *C*_*i*_ its impedance. The output of the inhibitory synapse into the dendrite or soma is a negative current pulse, the inhibitory postsynaptic current (IPSC).

### Excitatory synapse mapping

This subsection examines the functioning of an excitatory synapse, including the roles of calcium ions, glutamate, AMPARs, NMDARs, synaptic plasticity, and the translation of these biological processes into an equivalent electric-circuit model.

The function of an excitatory synapse (fig. 5) is similar to that of an inhibitory synapse, but the plasticity associated with spines adds complexity to the model. After lowpass filtering, excitatory input pulses increase the postsynaptic membrane potential, and the synapse’s gain is modified depending on the input’s magnitude and the current membrane potential.

**Figure 5:**
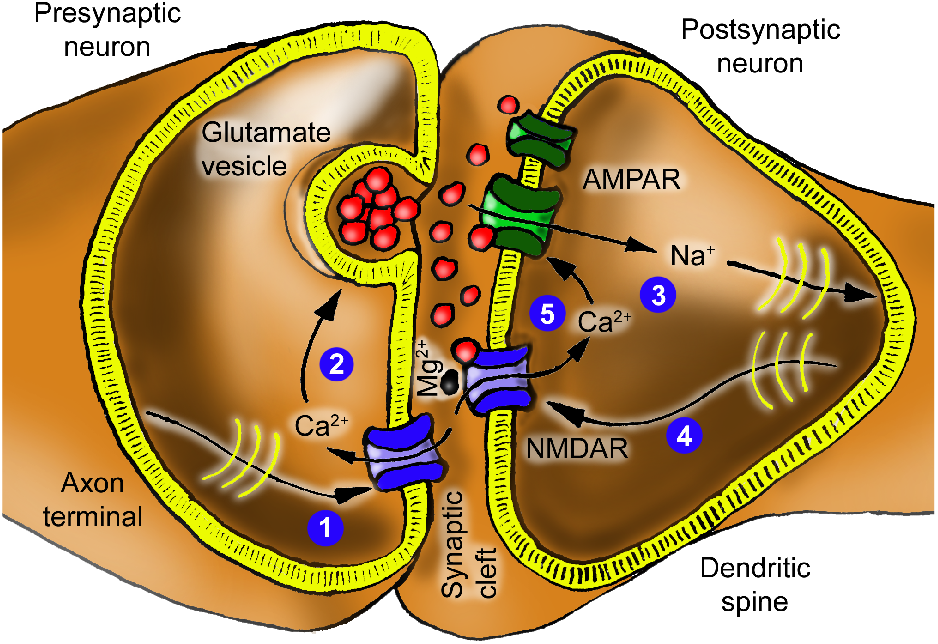
Excitatory synapse schematic. An action potential arriving at (1) causes the release of glutamate at (2), which then activates AMPARs, allowing a cation current at (3), here represented by sodium ions. This current travels to the soma and proximal dendrites, where it is lowpass filtered and fed back as the membrane voltage (4). At (5), this voltage and glutamate gate the NMDAR. Experiments have demonstrated the activity-dependence of the synaptic cleft’s calcium concentration [Ca^2+^]_e_ [6, 8].

In more detail, the arriving action potential enables calcium ions to enter the presynaptic terminal (1) and trigger the release of the neurotransmitter glutamate (2). Glutamate binds to AMPARs on the postsynaptic neuron, opening the channels to positively charged sodium and potassium ions (3) depolarizing the membrane potential.

The model opts for simplicity by depicting sodium ions only.

In addition, glutamate affects NMDA receptors involved in the neuron’s plasticity. The NMDA receptor is distinguished by its gating mechanism [31, 70]. While the binding of glutamate is essential, it alone is insufficient to open the channel. A magnesium ion is a gatekeeper that blocks the channel in a graded relation to the neuron’s membrane potential [51, 44]. Depolarization of the neuron removes this magnesium block (4), enabling calcium to flow through the NMDA receptor channel (5). This calcium influx regulates *the number* of AMPA receptors constituting the synaptic weight through a cascade of downstream reactions [52, 31].

The source of calcium ions is the synaptic cleft. This calcium is consumed *both* by the presynaptic terminal via the Ca.V channel *and* the NMDA channels via the “calcium path” illustrated in Fig. 6, which shows an electric-circuit equivalent for the excitatory synapse. Again, this is a direct translation of the biochemical processes of the excitatory synapse in fig. 5. Similar to the “Cl-” voltage source in the inhibitory synapse circuit, the “Ca2+” and “Na+” voltage sources represent the differences between the resting potential and the equilibrium potentials for calcium and sodium, respectively.

**Figure 6:**
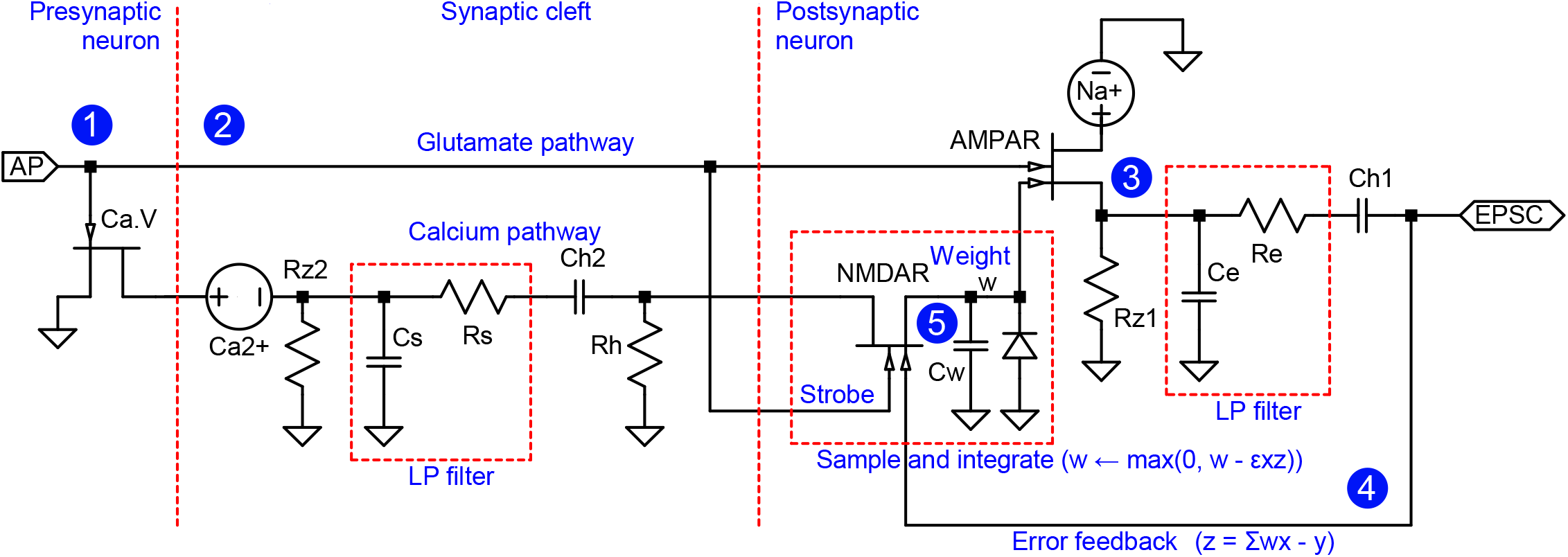
Excitatory-synapse circuit equivalent. In analogy with fig. 4, the above circuit models the excitatory synapse described in fig. 5 by representing the flow of neurotransmitters as electrical currents. Transistors represent gated ion channels. Locations indicated by circled numbers 1-5 correspond to identically marked locations in fig. 5. An action potential arriving at (1) causes the release of glutamate at (2), which then activates AMPARs, allowing a sodium current at (3). This current travels to the proximal compartment, where it is lowpass filtered and fed back as the membrane voltage (4). At (5), this voltage and glutamate gate the NMDAR transistor. Together with the capacitor *C*_*w*_, it represents the downstream cascade, eventually integrating the signal into the number of AMPARs. The annotations clarify the function of the components and indicate the adaptive filter function in analogy with fig. 7. The calcium pathway and the sample-and-integrate block are essential to the model, as NMDAR-mediated calcium influx regulates the number of AMPARs and thereby defines the synaptic weight. The capacitors *C*_*h*1_ and *C*_*h*2_ and the resistor *R*_*h*_ represent the cellular machinery retaining homeostatic equilibrium. The function of the NMDAR is based on the deviations from this equilibrium (“AC-analysis”).

A voltage pulse (1), representing the presynaptic action potential and corresponding glutamate release (2), gates injection of a positive current through the AMPAR. This current (3) is lowpass filtered as it travels to the soma, with a cut-off frequency of *f*_*c*_ = 1*/*(2*πR*_*e*_*C*_*e*_), which can vary significantly between different synapses within the same neuron. The filtering characteristics of excitatory synapses, much like those of inhibitory synapses, are influenced by their spatial positions and can be modeled using the *R*_*e*_ and *C*_*e*_ components. The culmination of this process is an excitatory postsynaptic current (EPSC) that is integrated with other EPSCs and inhibitory postsynaptic currents (IPSCs) by the membrane capacitance *C*_*m*_ of the soma and proximal dendrites. The ensuing (AC, alternating current) membrane potential *v*_*m*_ is electrotonically propagated throughout the cell (4).

The calcium concentration in the synaptic cleft is depleted when a presynaptic action pulse arrives, because the pulse causes calcium channels to open, consuming some of the synaptic cleft’s calcium content [6, 8]. This reduction of [Ca^2+^]_e_ upon activity in the synaptic cleft reduces the driving force for calcium entry through the NMDAR, which plays a crucial role in the mechanism underlying synaptic enhancement [4].

The cluster of NMDARs, modeled here as a dual-gate transistor, senses the membrane potential with one gate, whereas the other (“strobe”) recognizes glutamate activation, enabling synaptic weight modification. The synaptic cleft acts as a calcium buffer and effectively low-pass filters the calcium-encoded signal with a time constant given by *R*_*s*_*C*_*s*_. In this context, “buffer” refers to a reservoir with large—but finite—capacity. As a result, perturbations such as tapping into it still produce noticeable effects.

The voltage across the capacitor *C*_*w*_ represents the synaptic weight, which governs the variable number of AMPARs. In this model, the cluster of AMPARs is represented by a dual-gate transistor, enabling modulation by NMDARs. Since the number of AMPARs must be non-negative, the voltage across *C*_*w*_ is constrained to be non-negative. This constraint is enforced by a diode connected in parallel with *C*_*w*_.

The NMDARs’ key role is to act as a *multiplier* of deviations *x* and *z* from equilibrium in presynaptic activity and error feedback, respectively. The external calcium concentration [Ca^2+^]_e_ provides a lowpass-filtered copy of *x*, while voltage feedback from the soma supplies *z*. The product *xz* determines whether depression or potentiation occurs, depending on its sign.

*C*_*h*1_, *C*_*h*2_ and *R*_*h*_ represent homeostasis of the calcium pathway. Similar to the inhibitory synapse, they can also be seen as a compartmentalization of the model, which does not impair the signal transmission.

The complete equivalent circuit for the neuron can be formed by combining the circuits illustrated in fig. 4 and fig. 6. However, before proceeding to show that the complete circuit implements an adaptive filter, the next subsection offers a brief review of such filters.

### Internal structure and operation of an adaptive filter

This subsection concisely reviews the fundamental adaptive filter (fig. 7), which can be thought of as a procedure or algorithm. Its principal function is to find weights *w*_1_, *w*_2_, …, *w*_*n*_ such that the weighted sum Σ*w*_*k*_*x*_*k*_ of candidate or component signals *x*_1_, *x*_2_, …, *x*_*n*_ approximates a reference signal *y*. The component signals may originate from different sources or be derived from a single input *x* using a delay line or a filter bank as a signal decomposer.

**Figure 7:**
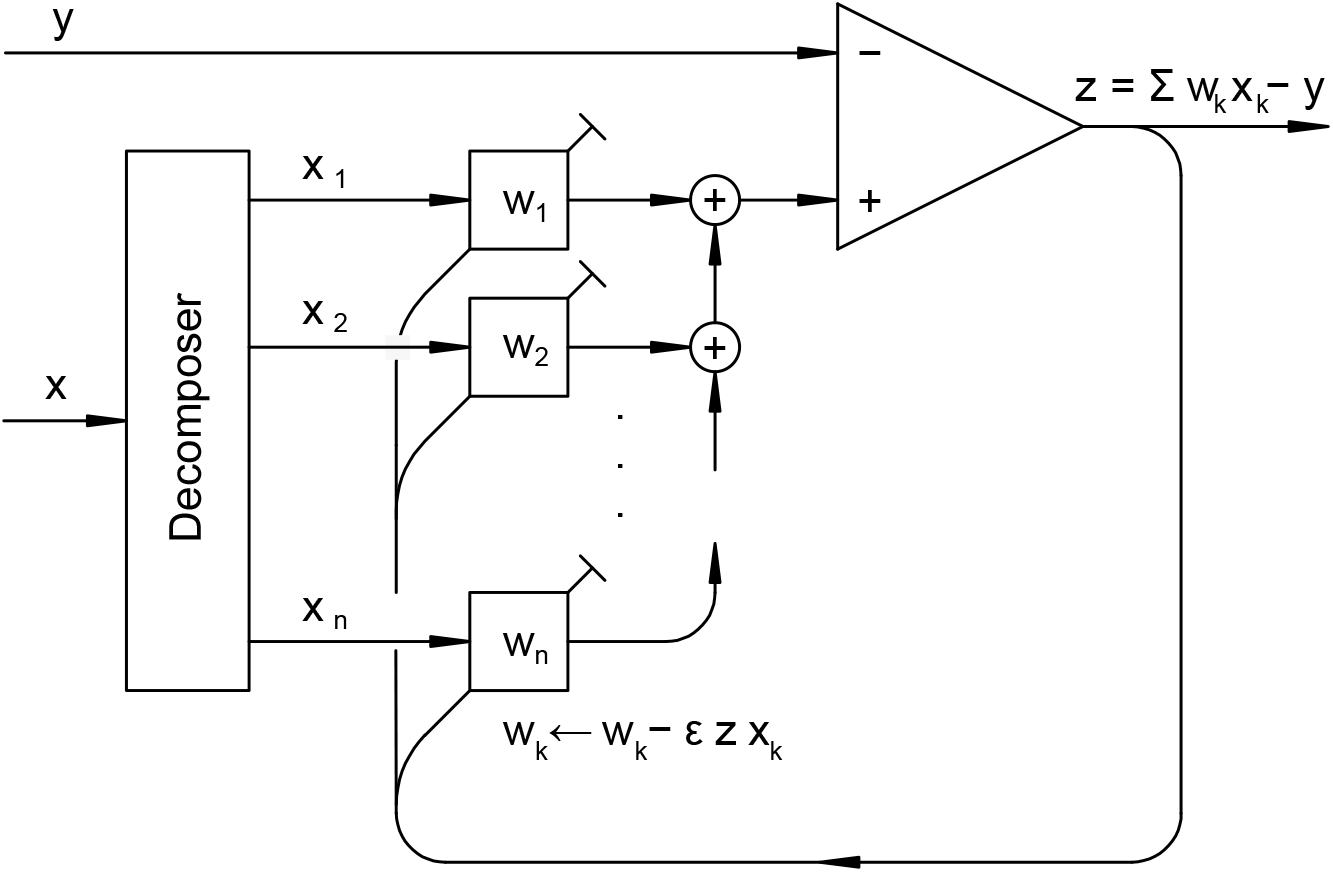
Structure of an adaptive filter [26]. A box with input *x*_*k*_ and tunable weight *w*_*k*_ computes the product *x*_*k*_*w*_*k*_ and corresponds to the excitatory synapse in fig. 6. The feedback *z* is crucial for adjusting the weights. The image is a graph representation of algorithm 1.

The adaptive filter can be interpreted in different ways, depending on one’s perspective. On the one hand, a biologist might see it as a system that maintains a balance between excitatory and inhibitory inputs [36, 55, 14, 69]. Notably, this differs from homeostasis because the inhibitory-excitatory balance adjusts synaptic weights so that the current weighted excitatory inputs match the inhibitory inputs as well as possible. It is important to note that, although there are multiple inhibitory inputs, the inhibitory weights are fixed according to Gray’s rules.

Therefore, we can represent the weighted sum of all inhibitory inputs as a single scalar variable *y*.

On the other hand, a physicist might view the filter as performing a wavelet transform of the signal *y* using wavelets *x*_*k*_ [43], with the weights serving as transform coefficients.

Depending on how the filter is connected, it can perform a variety of essential signal processing tasks such as model creation, inverse model creation, prediction, and interference cancellation [27]. Particularly relevant for biological systems is a configuration suggested to address the sensorimotor association problem, or the process by which the brain learns which neuron is connected to which muscle [47].

#### Algorithm 1: The basic LMS algorithm.

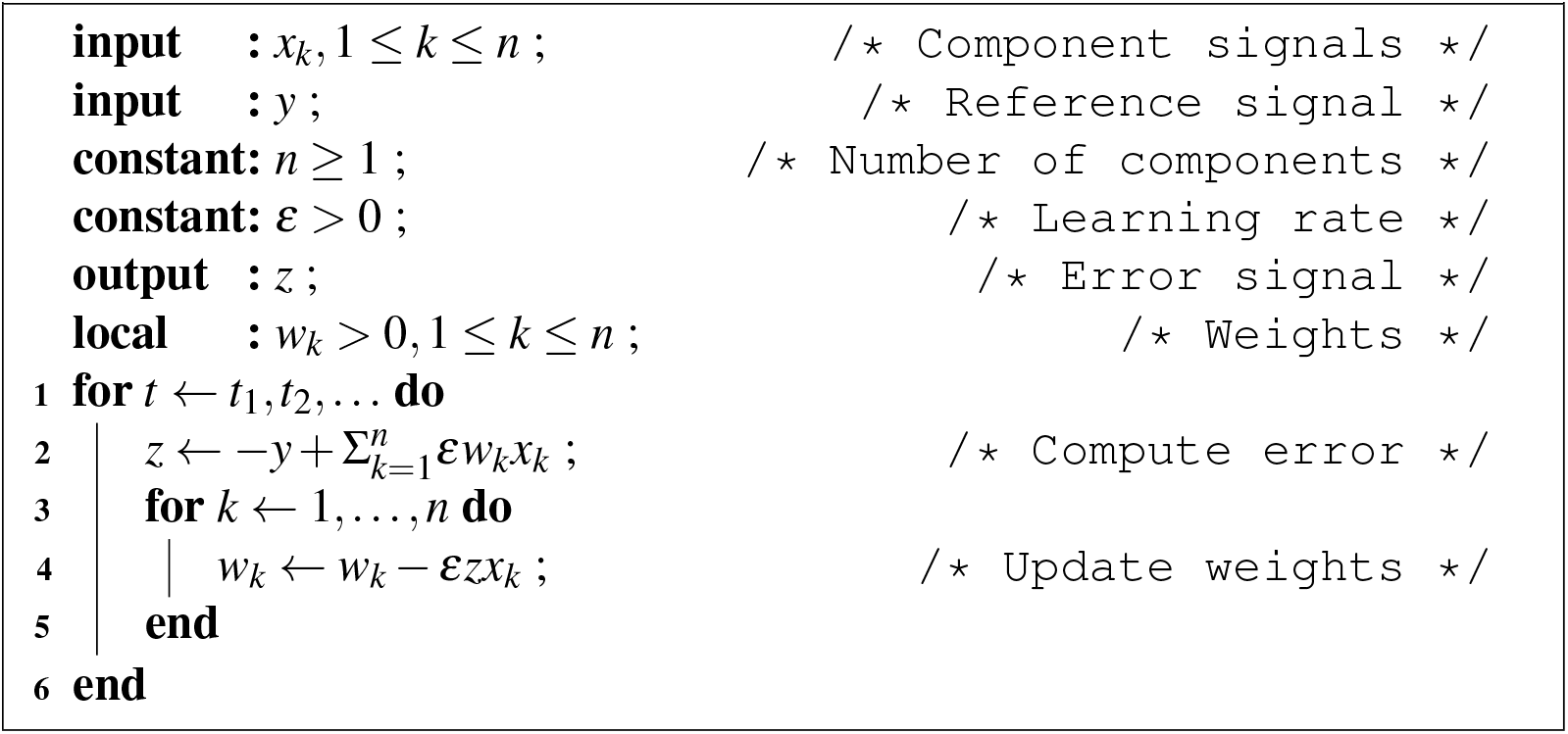

The Least Mean Squares (LMS) algorithm (algorithm 1) [26], also known as the Widrow-Hoff LMS rule, is a method for updating the weights of an adaptive filter. It operates in discrete time steps *t* = *t*_1_, *t*_2_, …, where at each step it calculates the error feedback *z*, which is the difference between the weighted sum of the input signals *x*_*k*_ and the reference signal *y*. Then, it updates all the weights *w*_*k*_ by subtracting the associated feedback corrections, which are calculated as Δ*w*_*k*_ = *ε zx*_*k*_, where *ε* is a learning rate. This learning rate is a positive constant, and its selection involves a balance between the convergence speed and stability against noise.

The convergence of the adaptive filter can be understood intuitively as follows: Suppose that some weight *w*_*j*_ is slightly too large and that the corresponding input *x* _*j*_ is positive. Then the error *z* will also tend to be positive and will be fed back to cause a reduction of the weight *w*_*j*_ by *εzx*_*j*_. A similar argument can be used when instead *w*_*j*_ is too small or *x* _*j*_ is negative. Proving the convergence of the weights formally can be difficult in general, but the LMS rule has proven to be robust in practical applications [27, 77].

### Understanding the neuron as an adaptive filter

Here, it is established that the neuron’s equivalent circuit operates as an adaptive filter, suggesting that the neuron also embodies this functionality.

#### The neuron’s equivalent circuit as an adaptive filter

Interpreting the neuron as an adaptive filter is greatly simplified by modeling the neuron as an equivalent electric circuit. The combination of the synapse circuits in fig. 4 and fig. 6 into a circuit equivalent for the neuron is shown in fig. 8. This circuit converts the spiketrain input to membrane potential. The subsequent output conversion of the membrane potential to an output spiketrain and the application of an activation function *ϕ*(*z*) are omitted here because a mechanistic model for them has been presented elsewhere [49] and does not directly influence the input conversion.

**Figure 8:**
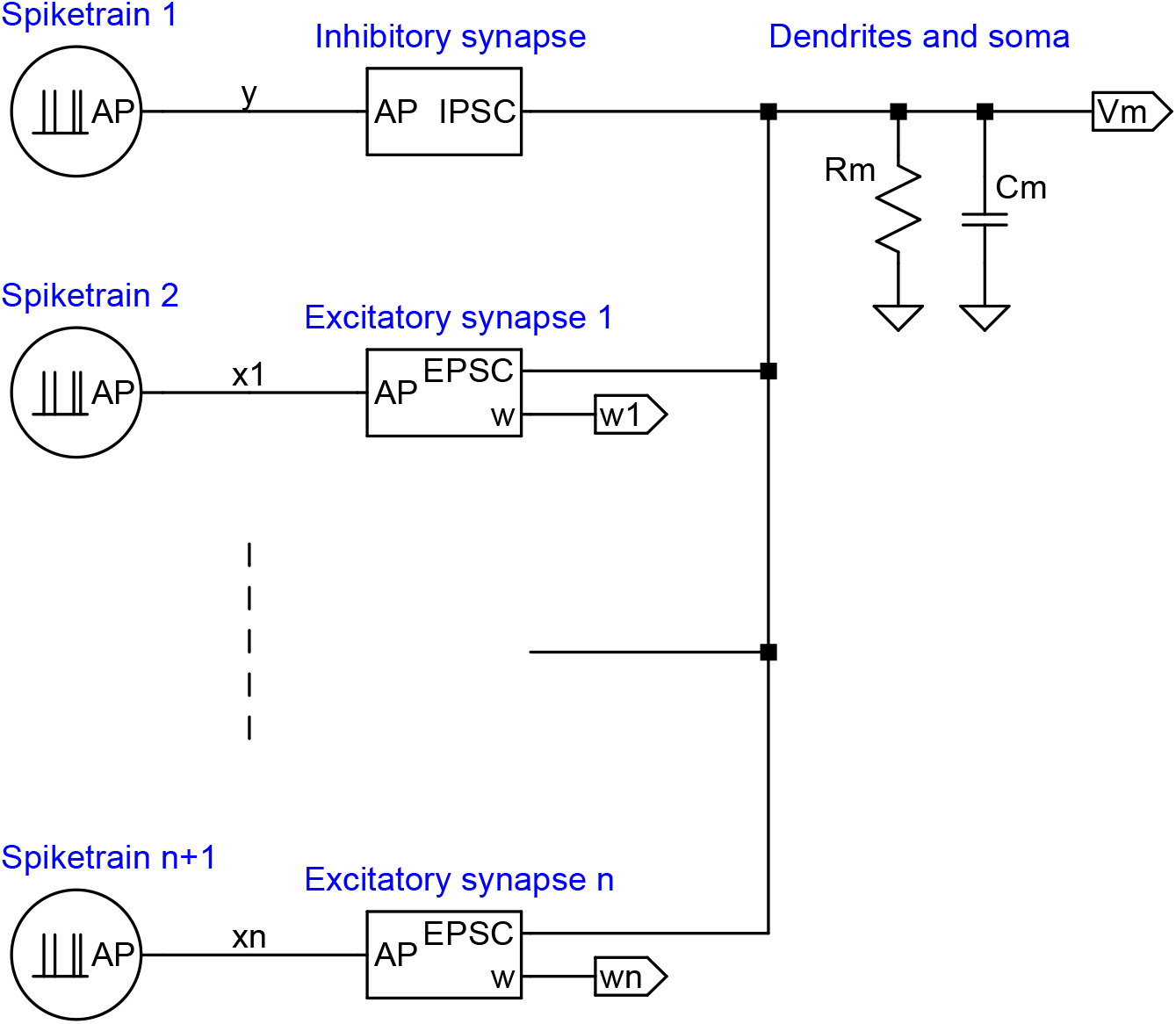
The neuron equivalent circuit. The IPSC and EPSC blocks are defined in fig. 4 and fig. 6, respectively. Labels *y, x*1,…, *xn* denote unfiltered action potential (spiketrain) inputs, which are lowpass filtered in the synapse blocks. The subsequent conversion of *v*_*m*_ back to an output spiketrain and application of an activation function are described elsewhere [49].

The side-by-side comparison of the adaptive filter, as shown in fig. 7 and the neuron model presented in fig. 8 offers detailed agreement, indicating that both the circuit and, by extension, the neuron implements a modified LMS algorithm (algorithm 2). The match between the circuit and the adaptive filter is corroborated below by illustrating how the circuit realizes the summation operations, error feedback, and weight updates. Furthermore, an explanation is provided for the scenario where component inputs are redundant or linearly dependent, a common condition for biological neurons.

##### Algorithm 2: The modified LMS algorithm, full neuron version with activation function included. The inputs are assumed to already be lowpass filtered.

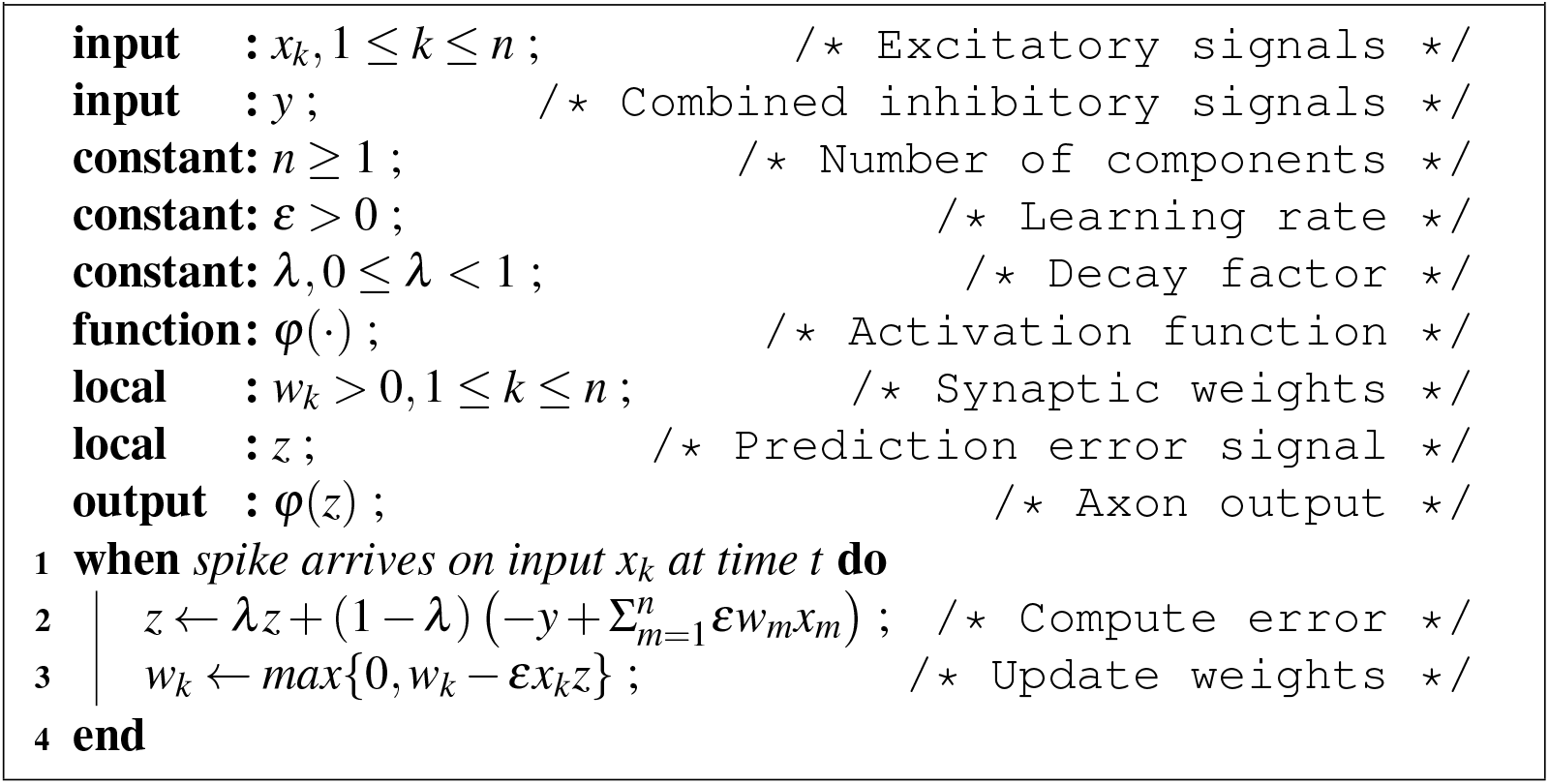

##### Summation operations

When comparing the functional blocks in fig. 7 with those in fig. 8, it is evident that the summation operations in the adaptive filter align with the addition of currents in the neuron’s equivalent circuit, as Kirchhoff’s law dictates. This law states that the sum of currents entering a junction must equal the sum of currents leaving it, mirroring the summation process in the adaptive filter.

##### Error feedback

A rapid error feedback signal, labeled by *z* in fig. 7 and fig. 6, is essential for the functioning of the adaptive filter, as is visible in algorithms 1 and 2. This feedback is provided by the membrane potential *v*_*m*_ created by the total of the IPSC and EPSC currents passing through the impedance consisting of the membrane resistance *R*_*m*_ in parallel with the membrane capacitance *C*_*m*_. The feedback signal accesses all synapses within the neuron via their connections to the soma. The lowpass filtering by *R*_*m*_*C*_*m*_ introduces a decay or “forget” factor *λ*, 0 ≤ *λ <* 1, on line 2 of algorithm 2, slightly generalizing upon algorithm 1, which would have *λ* = 0.

Rongala et al. [60] have proposed that the membrane capacitance and resistance function as a lowpass filter, stabilizing external feedback in recurrent neural networks. This function is equally applicable to single neurons with internal feedback. In the diagram in fig. 8, this lowpass filter is represented by *R*_*m*_*C*_*m*_, and its impact is encapsulated in the decay factor *λ*. Notably, this parameter is essential but was not included in the original formulation of LMS learning.

In the biological neuron, the propagation of the *z* signal is nearly instantaneous due to its electrotonic conduction through the cytosol. Nature has elegantly arranged for signal feed-forward by current and feedback by voltage, thereby avoiding interference between the two signals. Notably, the model does not require the postsynaptic neuron’s generation of an action potential to adjust synaptic weights. The membrane potential provides the feedback. This is important because otherwise, a neuron with zero synaptic weights would have difficulties leaving this state.

##### Weight updates

The adaptive filter updates its weights *w*_*k*_ by the product of the inputs *x*_*k*_ and the error feedback *z*. The update uses a clever trick that stands out when viewing the involved circuitry, *i*.*e*., the plasticity circuitry of the excitatory synapse in fig. 6. The weight *w* is a charge held by the capacitor *C*_*w*_. The product of the input *x* and error *z* should update this weight. However, whereas the error is readily available in the circuit as the membrane potential *v*_*m*_, the signal *x* on the glutamate pathway is PFM encoded and is unusable for the update in this form. Although lowpass filtered in the dendrite and soma, it is directly summed into the membrane potential and is unavailable separately. *Fortunately, a lowpass-filtered version of x is available as the calcium concentration* [Ca^2+^]_e_ *in the synaptic cleft*. Thanks to this additional copy of *x*, the NMDAR transistor in the circuit and the ion channel in the neuron can crucially “compute”—pass a charge proportional to—the weight update by multiplying the calcium concentration representing *x* with the membrane potential *v*_*m*_ representing *z*. Experiment 1, described below, validates the above process.

##### Redundant and linearly dependent candidate inputs

Composing a signal *x* into components in engineering contexts relies on techniques such as a bandpass-filter bank or a Fast Fourier Transform. These methods ensure orthogonality, or at least linear independence, of the components *x*_*k*_. This independence is a critical requirement to guarantee the uniqueness of the weights. As a contrast, such a systematic decomposition is unfeasible from a biological perspective, resulting in identical reference inputs possibly giving rise to different synaptic weights. In the case of redundant component inputs, weights will converge (settle) towards a linear subspace rather than a specific point. Correlated component inputs can slow the convergence of the original LMS algorithm. This is because weights are updated simultaneously, which may lead to overshooting and oscillations. Here, evolution has provided an elegant solution for neurons because each synapse is updated individually and asynchronously by its own glutamate strobe signal (fig. 6), demonstrated in experiment 2 (cf. the “for” statement in algorithm 1 with the “when” statement in algorithm 2).

#### Implications of the neuron operating as an adaptive filter

The neuron behaving as an adaptive filter allows us to address the three key concerns in the introduction: the process of memory storage and retrieval, the combination of Hebbian and homeostatic plasticity, and the establishment of a unifying rule for synaptic plasticity. The proposed solutions to these problems are presented in the results section below. More generally, the adaptive filter provides a valuable conceptual model for understanding neuron populations and facilitates a succinct mathematical representation of these [48].

The following subsection conducts a series of experiments that confirm the functioning of the circuit as an adaptive filter.

### Experiment design

Two sets of experiments were carried out to explore and validate model properties. In the first experiment, the stability and convergence of the model were examined. The neuron model comprised one inhibitory synapse and two excitatory synapses (*n* = 2 in fig. 8). The task of the circuit was to determine the weights *w*_1_ and *w*_2_ so that the weighted sum of spiketrains 2 and 3 corresponded to spiketrain 1. The inputs were Pulse Frequency Modulated (PFM) spiketrains, effectively inhomogeneous Poisson processes, modulated by sine waves with a modulation depth of 67% (fig. 9). The mean spike frequency was 100 Hz, meaning that the maximum and minimum were 33 Hz and 167 Hz, respectively. Sine waves with frequencies that are integer multiples of each other were chosen as stimuli because highly efficient signal processing methods exist to detect and separate such signals embedded in noise. The modulations for the first experiment were 1 Hz and 2 Hz for spiketrains 2 and 3, respectively. The reference input, spiketrain 1, began with a modulation of 1 Hz but switched to 2 Hz after 150 s, ensuring a large number of spike arrivals and NMDA activation episodes. The signal switching after 150 s demonstrates the circuit’s responsiveness and detects if convergence to a particular value is merely accidental.

**Figure 9:**
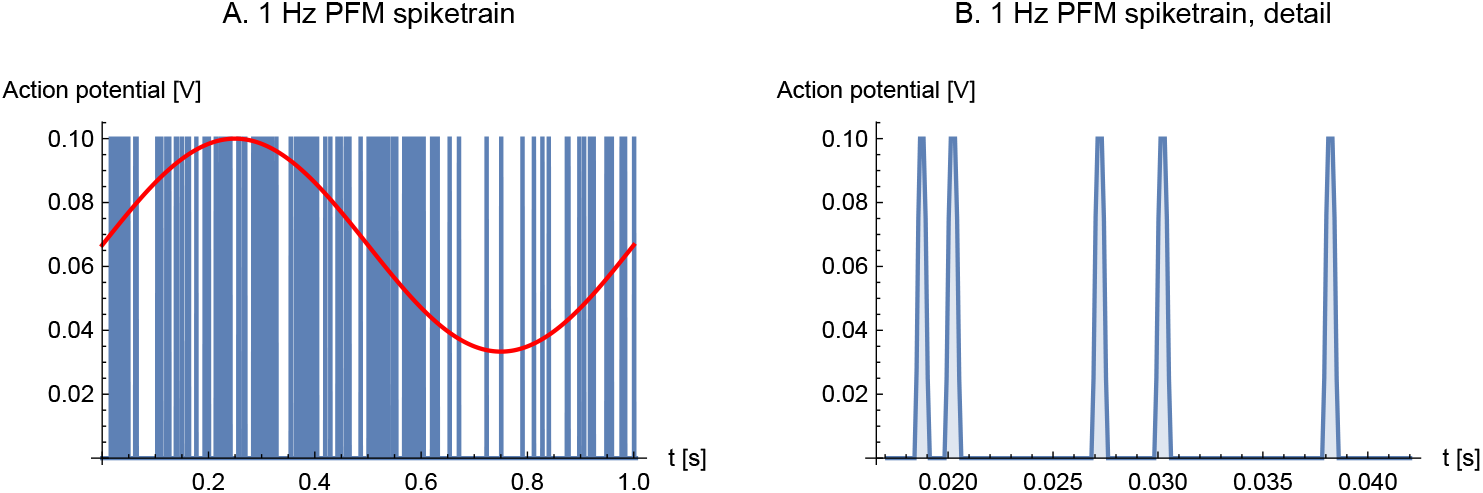
Input signals. A. Initial section of 1 Hz PFM spiketrain used as input. The overlaid sine wave shows the modulation signal. B. Detail. The spike width is 1 ms.

The second experiment aimed to study the model’s behavior in the presence of redundant input. In this experiment, a sine wave of 1 Hz modulated the inhibitory input, and a wave of 2 Hz modulated the first excitatory input (*x*_1_). The remaining five excitatory synapses *x*_2_, … *x*_6_ (*n* = 6 in fig. 8) received redundant input. During the first run, these inputs were synchronized, receiving the same spiketrain modulated at 1 Hz. In the second run, spiketrains 3-7 were modulated by 1 Hz but generated independently, mimicking the behavior of biological neurons, making them asynchronous, *i*.*e*., spikes arriving independently even though representing the same sine wave.

The component values used in these experiments are provided in table 1, and they roughly align with physiological values [28]. The *γ* parameters can be measured indirectly by their influence on the speed of adaptation and other time constants. In particular, the *γ*_NMDAR_ parameter is a lumped parameter that can be tuned to adjust the learning range *ε* over a wide range. The gamma parameters control the sensitivities of ion channels to gating parameters, such as neurotransmitter concentration. The primary gamma parameter *γ*_NMDAR_ directly controls calcium influx and thus learning speed. By adjusting this parameter, the neuron can regulate learning, setting it to zero to stop learning or to a high value for rapid learning. This parameter is likely to vary significantly depending on the state and type of neuron. While small calcium currents or gamma values would not cause unphysiological behavior, large values might. However, the experiments demonstrate that the circuit remains stable and functional even with substantial gamma values.

**Table 1:**
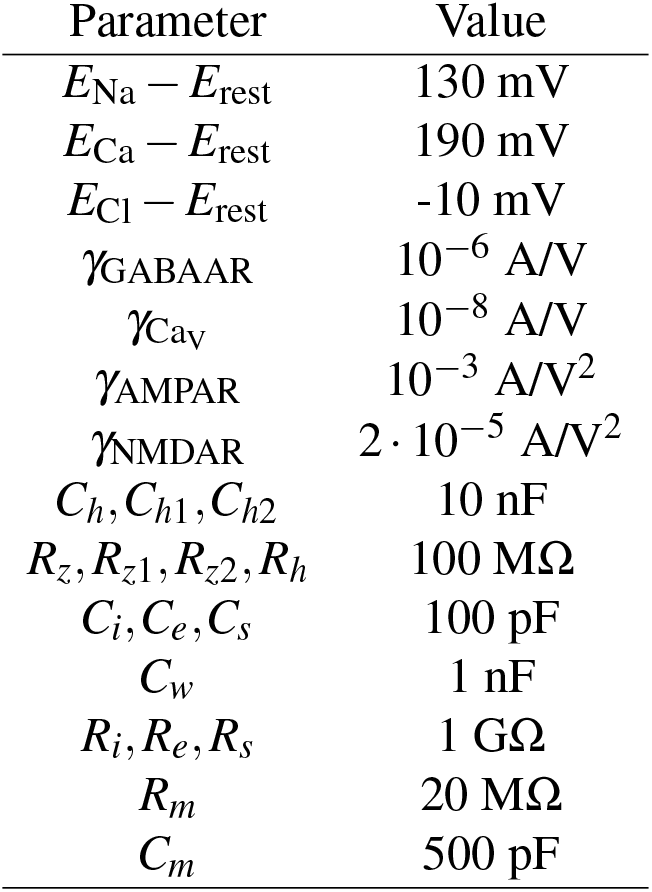
Model parameters. The *γ* parameters denote ion channel (transistor) gains.

Electrotonic signal propagation is assumed rapid over distances, such as the dendritic tree, while adhering to the RC constraints of the transmission path. Signal processing theory dictates that the feedback loop delay must be significantly shorter than the period of the maximum frequency transmitted through the circuit to maintain stability. Although other mechanisms could be involved in distal signal transmission, their presence is neither evident nor necessary for a mechanistic explanation.

The reversal potential for chloride ions is close to the resting potential at the inhibitory synapse, leading to a small electrochemical driving force for chloride ions, but this poses no issue, as it can be compensated for by a higher *γ*_GABAAR_ gain. Learning is typically considerably slower in biological neurons, but regardless, the circuit is robust and not overly sensitive to parameter variations. The experiments were conducted using the LTspice electronic-circuit simulator [41, 16].

## Results

### Experiment results

The first experiment demonstrates the convergence of weights *w*_1_ and *w*_2_. Initially, with the inhibitory input signal *y* modulated by a sine wave of 1 Hz, the ratio *w*_2_*/w*_1_ approaches zero as it should. This is because the input signal *x*_1_ is also modulated by a sine wave of 1 Hz, coinciding with the reference input, while the input signal *x*_2_ is modulated by a sine wave of 2 Hz, which is orthogonal to *y*. However, after 150 s, the modulation of *y* changes to 2 Hz, which instead coincides with the input signal *x*_2_. This time, the inverse ratio *w*_1_*/w*_2_ approaches zero. Fig. 10A depicts this convergence for two different values of NMDAR gain *γ*_NMDAR_. Low and high gain correspond to 2 *·* 10^−5^A/*V* ^2^ and 5 *·* 10^−5^A/*V* ^2^, respectively. The diagram shows that the circuit strives to enhance the weight of the excitatory input that aligns in frequency with the inhibitory input, while concurrently decreasing the weight of the other excitatory input that doesn’t match in frequency.

**Figure 10:**
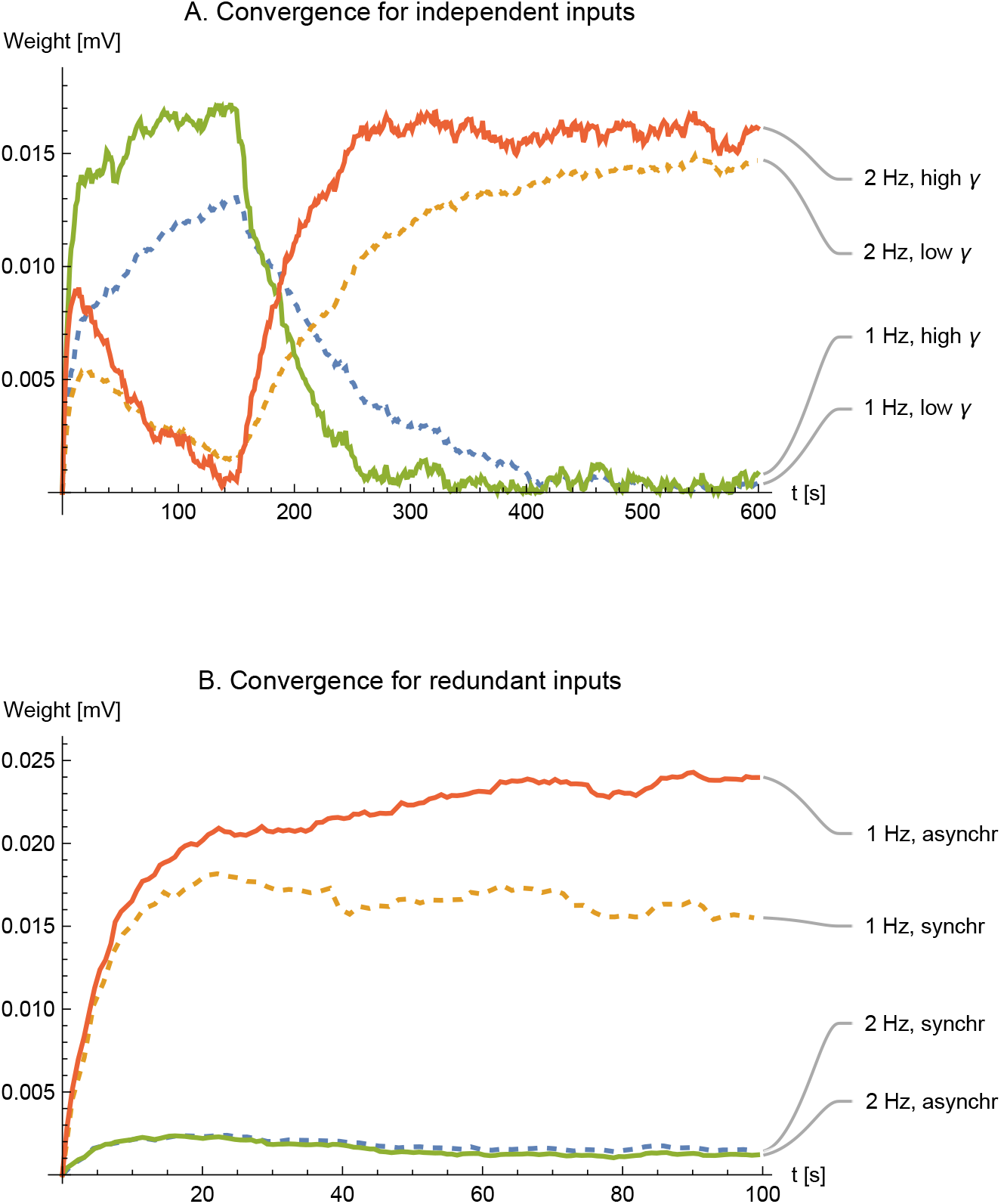
Convergence and stability of weights. **A**. Simulations show the convergence of the weights for two different values of NMDAR gain *γ* (dashed trace for *γ* = 2 *·* 10^*™*5^ *AV* ^*™*2^ and solid for *γ* = 5 *·* 10^*™*5^ *AV* ^*™*2^). Modulation of the reference signal changed from 1 Hz to 2 Hz at *t* = 150 s. **B**. Convergence for redundant inputs. The upper two traces show the sum *w*_2_ + *w*_3_ + … + *w*_6_, whereas the lower traces show *w*_1_. Weights cannot be negative. Dashed and solid traces are shown for synchronous and asynchronous spiketrains, respectively.

The second experiment shows what happens for multiple redundant excitatory inputs. In computer implementations of adaptive filters, simultaneous switching of redundant inputs can cause instability at high adaptation rates because of over-compensation. In biological neurons, action potentials typically arrive at different synapses asynchronously. Despite that, the experiments show that instability does not occur easily, even for synchronous arrivals.

In the first case, all the excitatory inputs are identical, so all strobe pulses are synchronous (dashed traces in fig. 10B). In the second case, the same sine wave of 2 Hz modulates the excitatory inputs, but otherwise, they are independent, so the strobe pulses are asynchronous (solid traces). The experiment shows faster convergence for asynchronous strobes.

Fig. 11 shows the evolution of synaptic cleft calcium concentration, calcium flow, IPSC, and EPSC during the first ten seconds of experiment 1. The bottom traces in panel A represent the [Ca^2+^]_*e*_ at the left terminal of *R*_*s*_ in the synaptic cleft for the two excitatory synapses. Despite some noise, the sine wave modulation of the input signals is evident, and the decrease in synaptic cleft calcium due to presynaptic activity is clearly visible. The top traces show the calcium flow into the NMDARs, where the amplitude of current variations decreases as the cell adjusts synaptic weights to balance inhibitory and excitatory inputs. Panel B illustrates the IPSC, reflecting the 1 Hz modulated input signal, while the EPSCs gradually increase from zero to counterbalance the larger IPSC.

**Figure 11:**
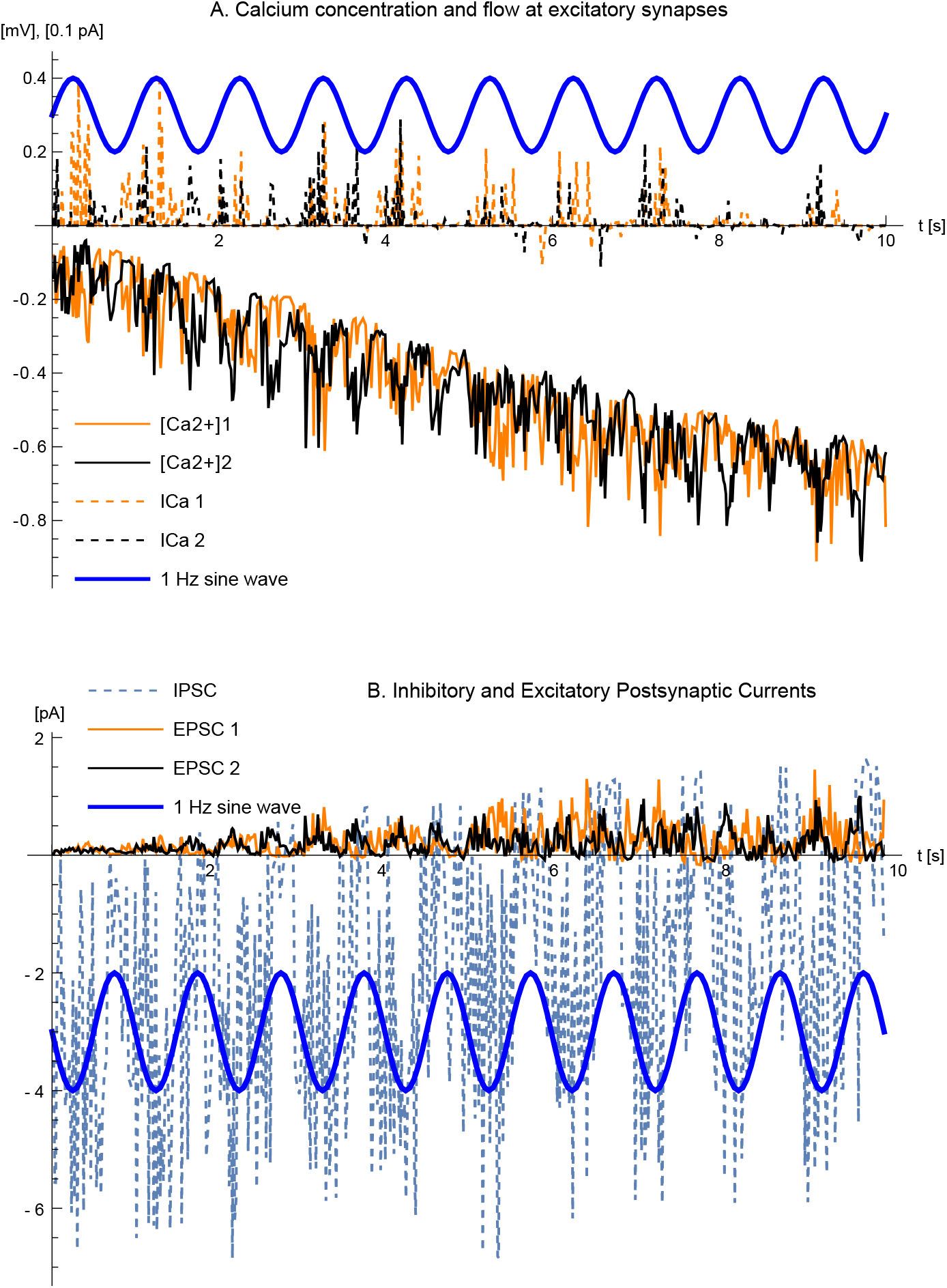
Calcium concentration, flow, IPSC, and EPSC. The figure illustrates the evolution during the first 10 seconds of experiment 1. **A**. Representations of [Ca^2+^]_*e*_ in the synaptic cleft (lower two traces) and Ca^2+^ flow through the NMDARs. **B**. IPSC and EPSC for the inhibitory and the two excitatory synapses. The blue thick traces are 1-Hz sine waves shown for reference only. The time axes of the panels are aligned.

It should be noted that understanding the entire neuron’s behavior based on individual currents is challenging, as its function involves feedback and relies on the cumulative effects of many small currents over time. It is easier to grasp the cell’s behavior through more abstract representations, such as the circuit equivalent in fig. 6, or at the algorithmic level in algorithms 1-2. This is a key takeaway of this article.

### Solutions to the three specific problems considered

This paper has suggested that a neuron functions and can be conceptualized as an adaptive filter with internal feedback. Such a neuron model enables straightforward solutions, presented below, to the three problems posed in the introduction.

#### 1. How does the neuron manage memory?

In this adaptive-filter model, memories are stored as synaptic weights. More precisely speaking, if the information is input in the form of an inhibitory signal *y*(*t*), the memory is formed in the neuron by adapting weights *w*_*k*_ so that *y* is balanced by the weighted sum Σ_*k*_*w*_*k*_*x*_*k*_ of the excitatory signals *x*_*k*_(*t*).

In principle, memory is subsequently retrieved whenever signals *y* and *x*_*k*_ are received by the neuron, and it computes and outputs the prediction error *z*, which relies on the weights. Alternatively, memory can be recalled by temporarily holding *y* at zero, whereby the neuron will output the approximation Σ_*k*_*w*_*k*_*x*_*k*_ ≈ *y* assembled by the same linear combination of excitatory signals.

#### 2. Is there a unifying synaptic learning rule?

The synaptic learning rule can be expressed as a variation of the Least Mean Squares (LMS) learning rule, modified to allow asynchronous weight updates, lowpass filtering of the feedback error, and the constraint that weights cannot be negative (cf. line 3 of algorithm 2):

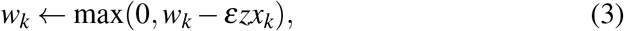

where *k* indicates synapse *k*, ***w*** = (*w*_1_, …, *w*_*n*_)^*T*^ is a vector denoting the numbers of AMPARs (synaptic weights), *z* represents the lowpass-filtered membrane potential *v*_*m*_ (error feedback), and the vector ***x*** = (*x*_1_, …, *x*_*n*_)^*T*^ signifies the vectors of local synaptic cleft calcium concentrations [Ca^2+^]_e_ (excitatory input). The learning rate *ε* depends on several biological parameters but is perhaps most directly controlled by the gain *γ* of the NMDAR. This rule is applicable for an arbitrary number of asynchronous inputs and is triggered on a spike arrival at excitatory input *k*.

#### 3. How do homeostatic and Hebbian plasticity balance?

The Hebbian-homeostatic balance emerges from the synaptic learning rule (3), inherently providing stability and subsuming both Hebbian and homeostatic plasticity. This learning rule attempts to minimize the mean square error between the desired output and the model’s prediction. It adjusts the synaptic weights based on the error signal which is the difference between the desired response and the actual output of the adaptive filter.

The stability of the modified LMS algorithm, irrespective of the sign of the input, comes from its inherent structure. The update rule is dependent on the product *zx*_*k*_ of the error *z* and the input *x*_*k*_. The multiplication *z · x*_*k*_ is directly implemented by the NMDAR. Even if the input changes sign, the direction of the weight update (whether to increase or decrease the weight) still appropriately aligns with the reduction of the overall error. This is because the error will also adjust its sign based on whether the prediction is above or below the desired outcome. Therefore, the product effectively guides the weight adjustments towards the direction that reduces the error, maintaining the stability of the learning process. Because the parameters *x*_*k*_ and *z* describe the signed deviations from the steady-state averages (homeostatic equilibria), the modified LMS rule offers automatic stabilization.

## Discussion

### The neuron as a differential element

The neuron uses membrane potential feedback during adaptation to adjust the excitatory synapse weights. This adjustment strives to balance inhibitory and excitatory input. Alternatively, this process can be described as the neuron’s attempt to predict the inhibitory input by excitatory input—the membrane potential encodes the *prediction error* [64]. Signal processing and control theory often refer to prediction error as the fundamental concept *innovation* [33]. It has frequently been discussed in neuroscience under different names, including *novelty* [37], *unexpectedness* [2], *decorrelation* [13], *surprise* [17], and *saliency* [74],

The critical operation for the plasticity of the neuron is the multiplication of the prediction error feedback *z*, represented by the membrane potential *v*_*m*_, with the excitatory input *x* available from the synaptic cleft external calcium concentration [Ca^2+^]_e_. Given the existence of this non-linear multiply mechanism, linear mechanisms can adjust a suitable homeostatic equilibrium or zero offset (*x*_0_, *z*_0_) by processes involving voltage-gated calcium channels (*zx*_0_) and metabotropic glutamate receptors (*z*_0_*x*).

A significant difference between a neuron and a classical adaptive filter is that the neuron’s weights cannot be negative. This is not a limitation because feeding a candidate signal *x* together with its negation (− x) achieves the same effect as a signed weight [12]. Incorporating such negations could be a function of the numerous local inhibitory neurons in the nervous system. Somewhat unexpectedly, this restriction to non-negative weights proves to be an advantage, as it enhances the expressive capabilities of neuron *populations* [48].

### Relation to neuromorphic engineering

Synapses have long been modeled as equations [61, 73], and particularly in the field of neuromorphic engineering, as electronic circuits [65]. These models are predominantly empirical, but they are typically too detailed in some respects and lack other crucial aspects to be useful for a mechanistic explanation of plasticity. This should not be construed as a dismissal of empirical models because they are significant in the development of neuroscience. Biologically inspired VLSI circuits are foundational, e.g., in achieving computational performance in neuromorphic engineering, and deserve recognition, even when biologically implausible. The specific concept that neuronal synapses function as lowpass filters, with an input spike typically resulting in a current shaped like an alpha function, has been a standard in neuron modeling. It has been systematically described by Gerstner and Kistler [20] but is not easy to attribute to any individual because it has evolved through cumulative research in the field.

Most circuit elements used here were introduced in neuron modeling well before the term “neuromorphic engineering” was coined in the late 1980s. The concept of the RC circuit as a foundation for neuron models was first proposed empirically by Lapicque in 1907 [38], and later, Cole and Curtis [9] developed it from a mechanistic perspective based on the bilayer structure of the membrane. Hodgkin and Huxley [29] suggested the existence of gated ion channels, which effectively functioned as transistors, although they were not depicted as such due to the unfamiliarity with transistors at the time.

Braeken et al. [7] constructed a FET transistor directly gated by glutamate. Dutta and Roy [15] introduced a variation of the Hodgkin-Huxley model where ion-sensitive FET transistors represent synapses. Gerstner and Kistler’s [20] description of the NMDA receptor aligns with the approach taken here, but their work is strictly confined to mathematical equations and does not involve introducing electronic components.

### Low-level model properties

Two salient features which distinguish the proposed model are the explicit dynamics of the synaptic cleft and the dual-purpose utilization of glutamate for both direct information transfer and as a strobe signal that facilitates weight adjustment. The necessity for a strobe input arises because if NMDARs were continuously active, weights would be diluted toward zero, resulting in information loss. It is crucial for plasticity that weights change only when there is meaningful input— that is, when activated by glutamate [31].

The circuit equivalent assumes that NMDARs operate at the same speed as AMPARs. In reality, NMDARs are slower and produce a burst of openings when triggered by glutamate, effectively performing a lowpass filtering. The model does not explicitly incorporate this property because the lowpass-filtered calcium input already accounts for the slowdown.

Several researchers have put forth adaptive filters as models for neuronal circuits in the cerebellum, utilizing *external* feedback [18, 78, 57]. Nevertheless, low-latency feedback is pivotal for the performance of an adaptive filter as it sets the maximum signal frequency content. External feedback is slower than internal feedback by several orders of magnitude (for pyramidal neurons, see, for example, [46, 1]).

The idea of a neuron functioning as a self-contained adaptive filter has been hypothesized [47, 42]. However, the model presented here appears to be the first wholly mechanistic model based exclusively on the known properties of ion channels.

While the chloride reversal potential acts as a limiter for large signal deflections and hyperpolarizations, this function is not crucial for mechanistically explaining plasticity when conducting a small-signal analysis. It is worth noting that the difference between the membrane potentials and the chloride reversal potential measured under physiological conditions *in vivo* tends to be greater than what the more common *in vitro* measurements suggest. More broadly speaking, in this model, the inhibitory synapse lacks plasticity and serves the simple role of signal inversion, lowpass filtering, and introducing an IPSC. A simple circuit can adequately model this functionality by a straightforward inverter followed by a lowpass filter, as detailed in equations (4)-(11) in the appendix.

The current paper follows Gray’s rules [23], which assert that excitatory synapses are linked to spines and exhibit neuroplasticity, while inhibitory synapses are considered static. Nevertheless, there are documented cases where inhibitory synapses demonstrate plasticity, although these do not involve spines or NMDA receptors. This means that the model proposed in this paper cannot be directly applied to such cases. A model aimed at understanding the plasticity of inhibitory synapses would need to be substantially different from the one presented here, even though similar analytical techniques using equivalent circuits could still be relevant.

For the studied GABAAR-AMPAR-NMDAR neurons, the model assumes that signals are conveyed by minor deviations from equilibrium. The general approach to model neurons by circuit equivalents can certainly also be applied in more general cases involving large deviations and steep changes in ion channel conductance depending on the operating conditions of the neuron, but in such cases, it will most likely be harder to find as simple an abstractions as (3).

### High-level model properties

Most neuronal plasticity experiments seem to apply uniform stimuli to both inhibitory and excitatory inputs. However, this study suggests that these inputs should be treated differently, as synaptic weight changes are heavily influenced by the relationship between them. Differentiating the stimuli for inhibitory and excitatory inputs is likely one of the most significant experimental proposals arising from this study.

A central prediction of the model is that the learning rate *ε*, or metaplasticity parameter, is directly related to the gain of the external-calcium-to-AMPAR cascade reflected by the lumped parameter *γ*_NMDAR_. Two interrelated experiments on real neurons could test this prediction. The first would test whether such a parameter is conceivable, e.g., by modifying the most convenient and accessible factor influencing the learning rate. The second would more exhaustively attempt to identify the factors affecting the gain and their interrelations.

One likely candidate for influencing the metaplasticity parameter is the baseline concentration of external calcium. Research, including a study using knockout mice [50], suggests that astrocytes regulate this concentration, significantly impacting LTP and LTD. Conveniently for experimentation, other studies have demonstrated that astrocyte activity can be modulated by noradrenaline [75], providing a potential experimental pathway for further investigation.

Conducting a sensitivity analysis to measure the factors influencing the learning rate is challenging because the above lumped-parameter gain of the NMDA receptor summarizes this sensitivity. Many factors influence this parameter, providing neurons with multiple adjustment methods. This adaptability is advantageous for the neuron, as it can select the most beneficial adjustment method. However, this complexity and the compensatory nature of these factors result in a broad operating range for each factor, making it hard to pinpoint parameter values.

When interpreting the circuits in fig. 4 and fig. 6 from an electrical engineering perspective, it appears that evolution has crafted a robust and minimalist solution. From a pure signal processing standpoint, the stability of neuronal functions strongly suggests the existence of feedback. The loop delay in this feedback must be short, pointing towards electrotonic propagation. The membrane potential is the sole feasible choice because the output spikes are too infrequent to provide swift feedback.

The neuron seems to use the biochemical equivalent of alternating current (AC) signals for communication, while the direct current (DC) level is regulated by homeostasis to maintain a suitable metabolic balance. It is hard to imagine a more efficient configuration of components capable of performing such a complex signal-processing task. Evolution has produced an elegant solution, utilizing current summation for feed-forward processes and voltage for feedback. The dual role of the glutamate pulse, acting both as a pulse-frequency modulated input and a strobe, is particularly striking.

Widrow and Hoff [76] initially introduced the abstract, high-level neuron model ADALINE (for ADAptive LInear NEuron), drawing inspiration from the McCulloch-Pitts neuron model [45]. This work predates the experimental discovery of ion channels by several years. Regrettably, Widrow and Hoff eventually abandoned ADALINE as a neuron model. Nevertheless, it became the foundation of the adaptive filter, which experienced dramatic advancements within the signal processing domain.

The LMS learning rule is known under various names in different contexts. In the field of artificial neural networks, it is often referred to as the “delta rule,” whereas in statistical learning theory, as the “covariance rule” [66]. These names all refer to the same concept: an iterative method for adjusting the weights of a learning model to minimize the mean square error ‖*z*‖ between the model’s prediction, which is the weighted sum of *x*_*k*_, and the actual data *y*. The proposed model is a mechanistic explanation of a modified LMS or covariance rule with asynchronous updates, restricted to non-negative weights and including a decay factor. Other major self-stabilizing learning rules are the Bienenstock-Cooper-Munro (BCM) rule [3] and the Oja rule [53]. However, these rules are theoretical constructs and, to the best of the author’s knowledge, lack mechanistic explanations.

The proposed model, when compared to biological neurons, exhibits several characteristics typical of biological neurons but not commonly found in other neuron models, at least not mechanistic ones:

1. It possesses the ability to record time-variable functions.
2. The model can learn without risking instability. This and the previous feature align with two of the three fundamental properties we initially aimed to achieve, as outlined in the introduction.
3. The capacity to “bootstrap” from a state where all synapse weights are zero is difficult for neurons relying on output spikes for plasticity.

The presented model does not include the process by which the neuron converts the membrane potential into the output spiketrain, including the activation function, because this process has been comprehensively addressed in a recent publication, which mechanistically explains this output process [49]. The current paper completes the picture of the neuron by providing a mechanistic explanation of the input process—the conversion back to internal potential from spike trains, including the plasticity.

### Related forms of plasticity

There is a vast body of literature exploring the mechanisms behind LTP and LTD, with many models focusing on the fine details of biophysical processes underlying synaptic plasticity. These models often aim to capture the intricate biochemical pathways that mediate calcium entry and AMPAR recruitment, but despite the level of detail in these studies, none, to the best of the author’s knowledge, provide a unified, mechanistic explanation of the neuron’s plasticity as a whole. This is largely due to the fact that these models operate at an overly specific level, focusing on the minutiae of chemical pathways where consensus is still lacking.

What is generally accepted, on the other hand, is the fundamental relationship that increased calcium levels lead to an increase in the number of AMPARs. This is the abstraction level at which the current model operates. While simplified, it effectively captures the core dynamic between calcium influx and synaptic weight modulation. This straightforward relationship, as demonstrated, is sufficient for explaining the broader behavior of the neuron.

In contrast to more detailed models that focus on replicating the specific biochemical pathways involved in LTP and LTD, this model offers a higher-level perspective that provides a mechanistic explanation of the neuron’s learning process. By adhering to the principle of Occam’s razor, the model demonstrates that a simpler, more abstract approach can sufficiently describe synaptic plasticity without the need for excessive complexity.

Several computational studies of synaptic plasticity acknowledge the importance of calcium current through NMDARs [67, 59, 22]. These models tend to focus heavily on spike-timing-dependent plasticity (STDP) and overlook the role of external calcium concentration, which complicates the acquisition of presynaptic activity [21].

It has been shown experimentally that STDP is not required for plasticity [32, 39], though it remains compatible with the model. A postsynaptic spike generates backward-propagating fluctuations in the membrane potential. While often called a backward-propagating action potential (BPAP), this spike is heavily lowpass filtered, appearing distally as a depolarization followed by hyperpolarization. If this “backwash” coincides with presynaptic activity, it can lead to an increase or decrease in synaptic weight, depending on the relative timing, as it contributes to the voltage error feedback, but a full analysis of this effect is beyond the scope of this paper and would require separate research.

Several other neuronal features have been discussed and speculatively related to plasticity, including electrical effects of the spine neck [24], location [63], intraspine action potentials [56], and shunting of synaptic currents by simultaneously active synapses on a single spine [35]. As for spine neck effects that passive filters can characterize, they benefit the neuron by increasing the diversity of synapse filter characteristics. However, the proposed model is generally independent of exotic features. Standard features of ion channels are entirely satisfactory for explaining all aspects of the model. Neither are exotic features deleterious for the model, as it is robust against noise in its capacity as an adaptive filter.

## Conclusions

Neuroscience research in many fields depends on detailed mechanistic knowledge of how neurons decode, process, store, and encode information. Examples of such fields are neural implants, interoception, and artificial intelligence, but progress in these fields has struggled with empirical and oversimplified neuron models.

This paper provides a complete state-of-the-art mechanistic model of a neuron’s signal processing, including the plasticity, in the milliseconds-to-minutes range. The model explains at the ion channel level how neurons convert input spiketrains to internal potential, including the adjustments of their synaptic weights. Crucial components of the model are the inclusion of synaptic cleft dynamics, the arrangement of internal feedback, and the multiple functions of the glutamate neurotransmitter. It is shown that memory recording can be identified with the weight adjustments of an adaptive filter. The neuron strives to balance the inhibitory and excitatory inputs. After adaptation, it can be regarded as an inhibitory input predictor, delivering the prediction error as output.

The mechanistic abstraction of the neuron as an adaptive filter constitutes an essential link to the realm of conceptual spaces [19] interposed between the cognitive and biological levels. It reduces the need for spiking-level simulations and simplifies the understanding of large assemblies and networks of neurons, elaborated in-depth in [48].

## Data availability statement

The netlists and symbol files designed and used during the current study are available on Github at https://github.com/drnil/neuroplasticity.

## Author contributions

The single author is responsible for all aspects of this research.

## Acknowledgments

This research was funded in part by the European Commission FP7 project THE (“The Hand Embodied”) under grant agreement 248587. This material is also based upon work supported by the Air Force Office of Scientific Research under award number FA8655-25-1-7007. The remainder was covered by RISE Research Institutes of Sweden internal funding for exploratory research.

ChatGPT-4 [54] was used to proofread parts of the text using the prompt: “Please correct and improve the following passage.” The author verifies and takes full responsibility for this use of generative AI in the preparation of the manuscript.

## Appendix: Synapse circuit equations

The equations decribing the inhibitory synapse in fig. 4 are shown below for the purpose of illustrating the complexity of an equation-based approach. Positive directions for currents and voltages are down and to the right.

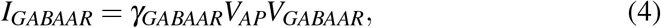

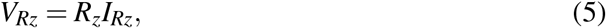

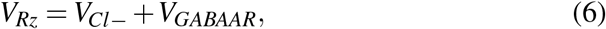

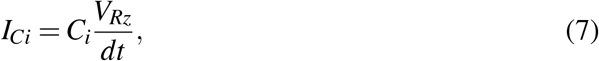

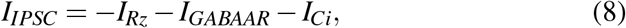

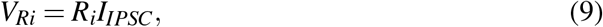

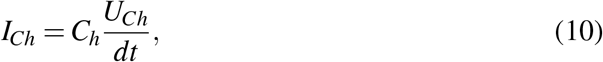

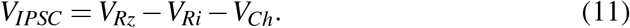

Equation (4) is the same as (2) in the section describing the inhibitory synapse.

The equations corresponding to the excitatory synapse in fig. 6 are

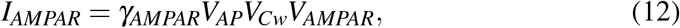

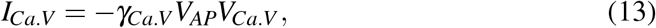

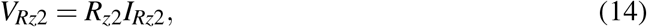

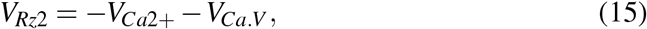

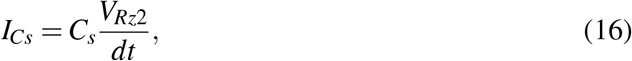

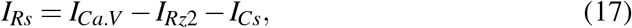

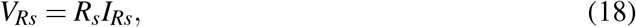

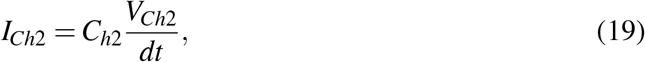

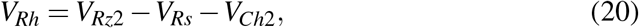

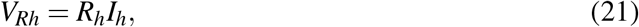

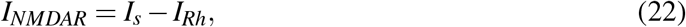

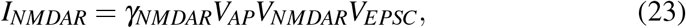

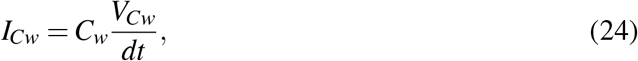

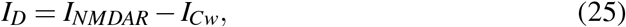

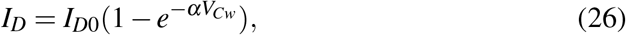

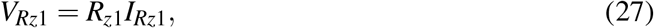

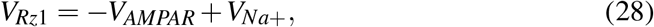

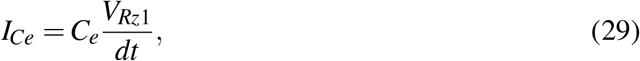

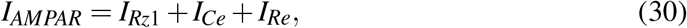

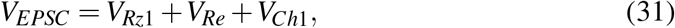

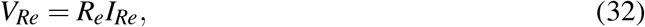

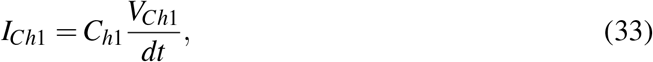

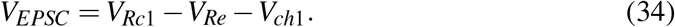

Complexity is introduced by the four non-linear equations (12), (13), (23), and (26), the five differential equations (16), (19), (24), (29), and (33), and the the feedback coupling by *V*_*EPSC*_ in (23).

The solution of the above equations, *i*.*e*., the model simulation, is carried out using the SPICE circuit simulator LTspice [41] via a netlist representation of the equivalent circuit. LTspice employs an Ordinary Differential Equation (ODE) solver with a modified trapezoidal integrator and a variable time step. It incorporates special numerical techniques for handling nonlinear equations, sparse matrices, and implicit integration [16].

The circuit simulator itself is agnostic with regard to neurons and neuroscience, which offers an advantage over specialized neuron simulators as there are no builtin hidden assumptions. The implementation is fully transparent, as all biological information is explicitly provided by the netlist.

Below, some details are provided how a system of non-linear ODEs is solved using the implicit trapezoidal method combined with Newton iterations.

The first step of the simulator is to express the network in the standard form

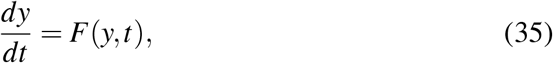

where *y* is a vector of the unknown currents and voltages. This is a straight-forward process accomplished by taking the time derivative of formulas which don’t already contain a time derivative. The implicit trapezoidal method then introduces difference approximations of the derivatives and updates the solution from *y*_*n*_ at time *t*_*n*_ to *y*_*n*+1_ at time *t*_*n*+1_ = *t*_*n*_ + Δ*t* using the formula

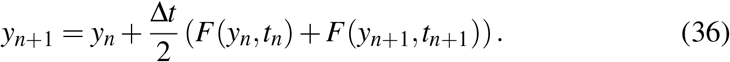

This equation is implicit because *y*_*n*+1_ appears on both sides of the equation, making it a non-linear algebraic equation when *F*(*y, t*) is non-linear. Here, Newton iterations are used to solve the implicit equation for *y*_*n*+1_. The equation to solve is

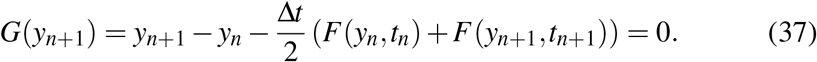

Newton iteration starts with an initial guess 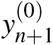 which is the previous time step’s solution, 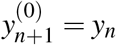. Iterative updates are then performed to refine the guess 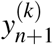 using the Newton iteration formula

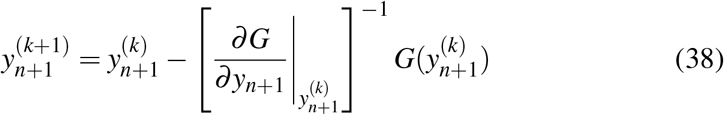

Here, 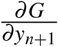 is the Jacobian of *G*(*y*_*n*+1_) with respect to *y*_*n*+1_. The current guess 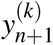 is the substituted into the equation for *G*(*y*_*n*+1_):

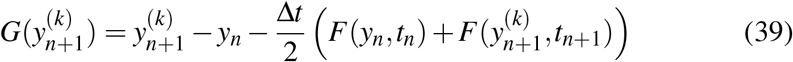

The Jacobian of *G*(*y*_*n*+1_) with respect to *y*_*n*+1_ is given by

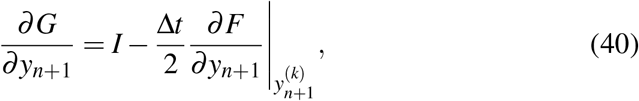

where *I* is the identity matrix and 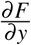 is the Jacobian matrix of *F*(*y, t*) with respect to *y*. After this, the Newton iteration formula computes the next guess,

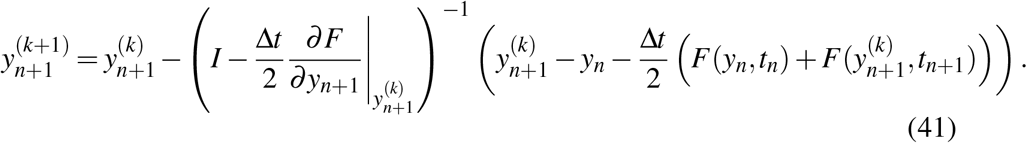

The iteration continues until the change in *y*_*n*+1_ between iterations is below a specified tolerance,

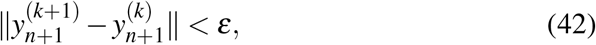

where *ε* is a small positive number.

